# Direct estimation of genome mutation rates from pedigrees in free-ranging baleen whales

**DOI:** 10.1101/2022.10.06.510775

**Authors:** Marcos Suárez-Menéndez, Martine Bérubé, Fabrício Furni, Vania E. Rivera-León, Mads-Peter Heide-Jørgensen, Finn Larsen, Richard Sears, Christian Ramp, Britas Klemens Eriksson, Rampal S. Etienne, Jooke Robbins, Per J. Palsbøll

**Affiliations:** Groningen Institute for Evolutionary Life Sciences, University of Groningen, Groningen, The Netherlands; Center for Coastal Studies, Provincetown, Massachusetts, United States of America; Greenland Institute of Natural Resources, Nuuk, Greenland; National Institute of Aquatic Resources, Kongens Lyngby, Denmark; Mingan Island Cetacean Study Inc., St. Lambert, Quebec, Canada; Scottish Oceans Institute, University of St. Andrews, St. Andrews, United Kingdom

## Abstract

Current low germline mutation rate (*μ*) estimates in baleen whales have greatly influenced research ranging from assessments of whaling impacts to evolutionary cancer biology. However, the reported rates were subject to methodological errors and uncertainty. We estimated *μ* directly from pedigrees in natural populations of four baleen whale species and the results were similar to primates. The implications of revised *μ* values include pre-exploitation population sizes at 14% of previous genetic diversity-based estimates and the conclusion that *μ* in itself is insufficient to explain low cancer rates in gigantic mammals (i.e., Peto’s Paradox). We demonstrate the feasibility of estimating *μ* from whole genome pedigree data in natural populations, which has wide-ranging implications for the many ecological and evolutionary inferences that rely on μ.

## Main

Adaptive evolution is ultimately driven by the appearance of novel genotypes, which in turn are a result of *de novo* germline mutations (*DNMs*) and recombination. The rate (*μ*, i.e., the probability of a nucleotide substitution per site per generation) of *DNMs* varies considerably among taxonomic groups. Multiple causes have been proposed to explain observed differences in *μ* (*1*), such as selection on *μ* itself, or a reflection of differences in physiological and cellular processes (e.g., metabolic rates, and DNA repair mechanisms). In addition, *μ* is a central parameter in evolution and population genetic inference and necessary when converting genetic estimates into more intuitive quantities, such as time and abundance. Most estimates of *μ* are obtained from DNA-based phylogenies (*μ*_PHY_) with dated branching points (e.g., from fossil records); an approach which is subject to multiple sources of error and uncertainty (*2*). However, the advent of inexpensive whole genome sequencing methodologies has enabled direct estimation of *μ* from pedigrees of individuals (*μ*_PED_), which relies on fewer assumptions and *μ*_PED_ is readily comparable among species providing an opportunity to test current hypotheses regarding the differences in *μ* and revisit key findings and hypotheses based on *μ*_PHY_.

Baleen whales (*Mysticeti, Cetacea*) include the largest and longest-lived mammals; aspects which have been invoked as the cause of low observed estimates of *μ*_PHY_ in this group of mammals. Their gigantic body size results in relatively low metabolic rates (compared to smaller-bodied mammals) and thus a lower expected *μ* (*3*). Indeed, estimates of *μ*_PHY_ in both the nuclear and mitochondrial (mt) genome of baleen whales were lower than similar estimates for smaller-bodied mammals, such as primates (*4*). The lower *μ*_PHY_ observed in baleen whales has, in turn, been hypothesized to explain low cancer rates in baleen whales, part of a phenomenon known as Peto’s Paradox (*5, 6*). However, the cause and consequences of such lower *μ*_PHY_ have been the subject of considerable debate (*4, 7*). Baleen whales were historically subjected to substantial human exploitation on a global scale from industrial whaling; but the full extent of that depletion remains unknown. Estimates of *μ*_PHY_ were applied to estimates of pre-exploitation abundance based on current levels of genetic diversity in some whale species (*8, 9*), yielding pre-exploitation abundance estimates 4-10 times higher than results obtained by other means (*10*). These high estimates of pre-exploitation whale abundance further imply that the historic oceans supported a much larger biomass of whales and prey.

Estimates of *μ*_PED_ in the nuclear genome have so far only been obtained from ~20 vertebrate species, mainly primates and model species raised in captivity (table S1). In contrast, estimates of *μ*_PED_ in natural populations are rare and so far confined to a single small group of individuals in small distinct population in only three species (a quartet of platypus, **Ornithorhynchus anatinus*, inhabiting the same creek, *11*, a septet in a single wolf, Canis lupus, pack, 12, and a known three-generation pedigree of collared flycatchers, Ficedula albicollis, in a study population, 13*). We demonstrate that *μ*_PED_ estimates are readily obtained by combining genetic relatedness analysis with genome sequencing in difficult-to-sample natural populations of non-model species without any prior pedigree knowledge.

We combined genetic parentage exclusion analysis (based on microsatellite genotypes) with molecular sex determination and mt DNA sequences to identify offspring and parents trios among free-ranging North Atlantic blue (*Balaenoptera musculus*), fin (*Balaenoptera physalus*), bowhead (*Balaena mysticetus*) and humpback (*Megaptera novaeangliae*) whales. Except for humpback whales, no prior pedigree information was available. However, despite a relatively low sampling probability in some species (e.g., ~4% for the bowhead whale, *14*), subsequent whole genome sequencing confirmed the initial identification of offspring and parent trios. Our estimate of *μ*_PED_ in baleen whales was based on *DNMs* detected in eight parent-offspring trios: one each in blue, fin and bowhead whales and five in humpback whales.

The complete genome was sequenced in all individuals to an average depth of ~30 and aligned against the blue whale genome (*15*). *DNMs* in the offspring genomes were identified by employing a modified version of Bergeron *et al.’s* (*16*) pipeline, manually curated and a subset of *DNMs* subsequently confirmed from Sanger sequencing. The accuracy of the modified pipeline and curation was assessed by reanalysing published data from macaques (*16, 17*), which yielded a similar estimate of *μ*. A total of 260 *DNMs* were identified in the eight offspring, among which 126 were phased, and 76% were paternal (tables S2 & S3). The majority of *DNMs* consisted of transitions from a strong to a weak base with a most likely non-functional impact (fig. S6). All biallelic sites in the putative offspring were consistent with the presumed parents, i.e., segregating according to Mendelian expectations. This was also the case for *DNMs* transmitted across two generations in humpback whales (Fig. 1). Age data for the sires were scarce, but suggested a positive correlation between the number of paternal *DNMs* and paternal age at conception, in agreement with other studies (*16, 18, 19*). No *DNMs* were shared between half-siblings (Fig. 1) suggesting that most *DNMs* arose during meiosis.

**Fig. 1.**
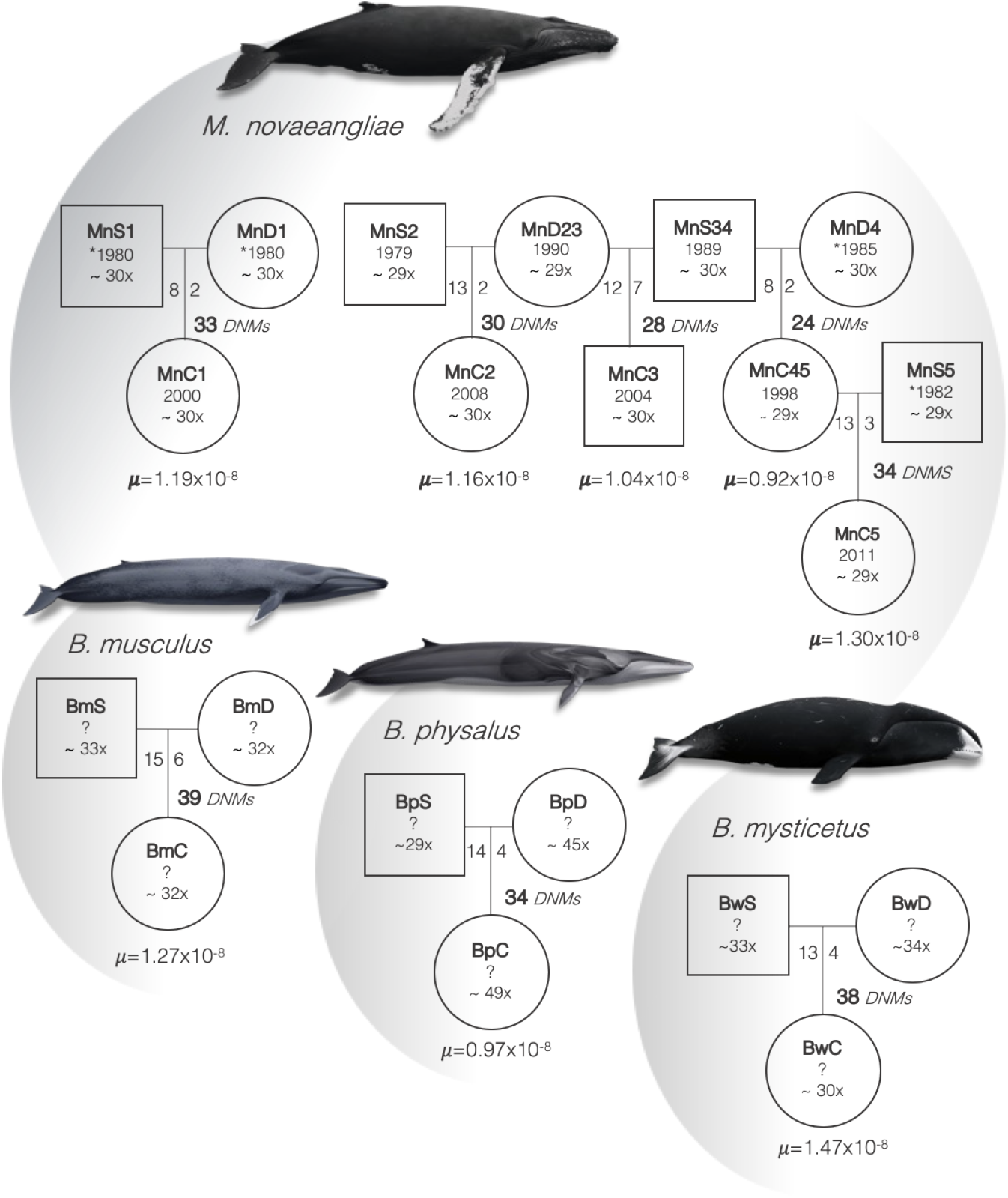
Summary of the estimation of *μ*_PED_ in the nuclear genome per parent-offspring trio in humpback, blue, bowhead and fin whales. Squares and circles denote males and females, respectively, and contain sample identification number; birth year (or the year in which the individual was first sighted, denoted by *), no previous sightings (denoted by ?) and mean genome sequence coverage. Along the vertical lines are listed; the total number of detected *DNMs* and the number of *DNMs* originating from the father (left) and the mother (right). *μ*_PED_: the estimate obtained from the specific trio. Whale illustrations by Frédérique Lucas (size of whales is not to scale).

*DNMs* were detected in 63% of the autosomal genome yielding estimates of *μ*_PED_ at 1.12×10^-8^ (range: .93×10^-8^-1.3×10^-8^) and 1.17×10^-8^ (range: .93×10^-8^-1.47×10^-8^) in both humpback whales alone and all baleen whales combined, respectively (Fig. 1, table S2). The latter estimate is very similar to estimates of *μ*_PED_ in humans and other large primates (Fig. 3, A) contrary to the notion of a lower *μ* in baleen whales. Prior acceptance of low *μ*_PHY_ estimates was mostly based on the hypothesized effects of a reduced metabolic rate in gigantic mammals (500× - 2,000× by weight relative to humans), resulting in lower levels of intracellular free radicals and consequently less DNA damage (*4*). It is possible, although somewhat implausible, that *μ* is lower in somatic cells compared to germline cells (*20*). However, in such a case, baleen whales would differ from other mammals in terms of a mechanism that solely lowers somatic, but not germline, mutation rates (e.g., DNA repair mechanisms). Barring such a unique mechanism, our findings do not provide sufficiently low rates of *μ* needed to explain Peto’s Paradox, i.e., lower rates of cancer in gigantic mammals (*7, 21*).

The hypervariable control region in the maternally transmitted mt genome is one of the most common genetic markers employed in population genetics towards a wide range of fundamental aspects in ecology and evolution. Contemporary levels of genetic diversity in baleen whales have been employed to infer unexpectedly high pre-exploitation abundances in North Atlantic baleen whales (*8*). These genetic assessments of the impacts of whaling were based on *μ*_PHY_. The much smaller size of the mitochondrial genome (16.5KB), implies few, if any, *DNMs* in each offspring. Accordingly, a large sample of mother-offspring pairs (*22*) is necessary to obtain an estimate of *μ*_PED_ in the mt genome. We capitalized on the long-term research on North Atlantic humpback whales (*Megaptera novaeangliae*) in the Gulf of Maine since the 1970s. We identified 141 maternal lineages among 848 humpback whales, characterized by 20 different mt haplotypes, each numbering between two and 49 whales and spanning one to four generations. *DNMs* in the mt control region were evident as heteroplasmy and detected in the oldest known female in nine maternal lineages and at minimum one descendant to confirm germline heteroplasmy. In total, heteroplasmy was detected in five, three and one lineages at sites 82, 258 and 235 (table S5), respectively. Heteroplasmy at site 82 was identified in three different mt haplotypes, confirming the presence of mutational hotspots (*23*). The *DNMs* detected as heteroplasmy in seven lineages, yielded mt control region haplotypes already present in the population (table S5), i.e., *DNMs* that may go unaccounted in estimates of *μ*_PHY_ (*9*).

We inferred *μ*_PED_ from the generational change in the frequency of each variant at the heteroplasmic site as per Hendy and colleagues (fig. 2, *24*) from massive parallel sequence data with an average read depth at 6,900 (*20*). In total, 47 maternal transmissions of a germline mt heteroplasmy were used to infer a median estimate of the number of segregating units at 10.4 (interquartile range, *IQR* 5-28.9, fig. S7) during the oocyte bottleneck. The number of segregating units was similar to, but at the lower end, of the range of values reported in other species (*22, 26–29*). The final estimate of *μ*_PED_ for the mt control region was at 4.3×10^-6^ (*IQR* 1.55×10^-6^-8.85×10^-6^, table S6), similar to comparable estimates of *μ*_PED_ for the same mt region in humans (3.8×10^-6^; *IQR* 1.18×10^-6^-1.16×10^-5^, *27*).

**Fig. 2.**
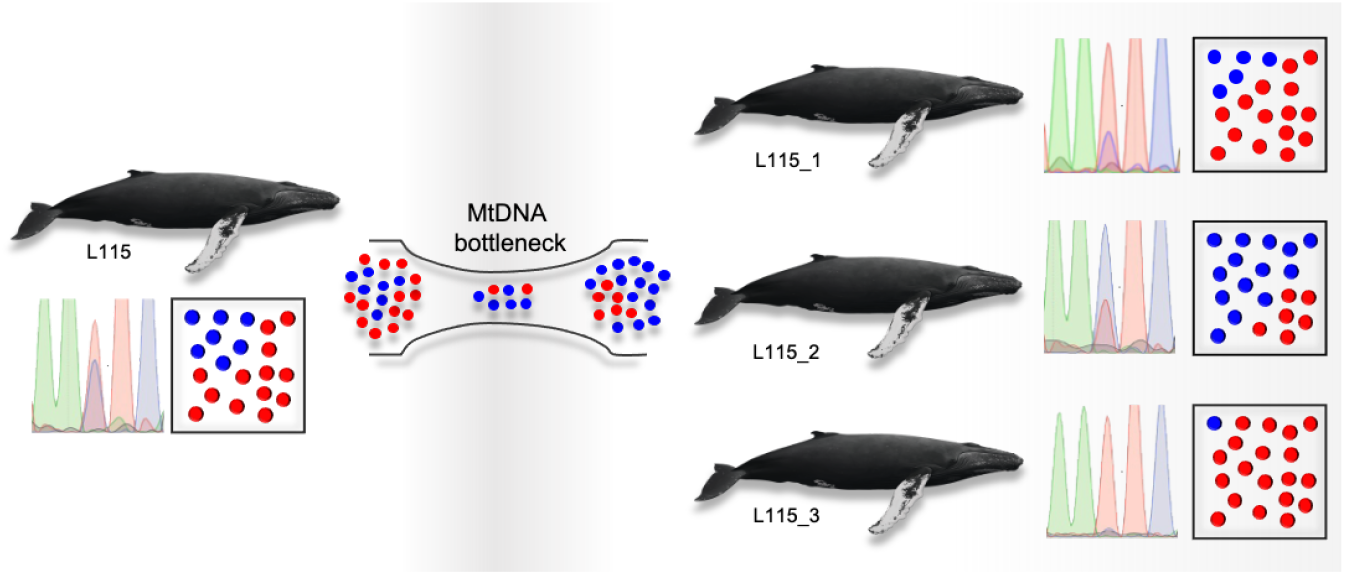
An example of the change in mt haplotype frequencies of a point heteroplasmy after the transmission bottleneck during oocyte production in a humpback whale cow and her three calves as evident in Sanger sequencing chromatograms (left-hand side square) and sequence read frequencies (right-hand side square). The relative frequencies of the two mt haplotypes were reversed in calf 2 compared to the mother. In calf 3, the heteroplasmy was undetectable by Sanger sequencing but detected among the sequenced reads. Whale illustrations by Frédérique Lucas.

We assessed the consequence of our findings on previous estimates of pre-exploitation abundance of North Atlantic humpback whales that were inferred from contemporary levels of genetic diversity (*8, 9, 30*). Insights into the pre-exploitation abundance of baleen whales are not only relevant to their current level of endangerment, but also provide fundamental insights into the overall state of the historic oceans and the biomass they supported prior to human overexploitation. Our re-analysis was conducted as in previous studies (*8, 30*) except for using the value of *μ*_PED_ for the mt control regions estimated here. We estimated an effective female population size of 2,561 (*IQR* 1,245-7,128), which translated into an abundance of 20,494 humpback whales (*IQR* 9,964-57,030). This estimate corresponds to only 14% of the lowest *μ*_PHY_-based estimate based on mt control region sequences (150,000, credited to *9*, by *30*). Notably, the abundance based on our estimate of *μ*_PED_, is similar to non-genetic estimates of pre-exploitation humpback whale abundance (17,151-22,647, *10*). Together these results suggest that estimates of pre-exploitation whale abundance based on *μ*_PHY_ are likely strongly upward biased. Consequently, the most recent estimate of abundance of North Atlantic humpback whales (10,752, *31*) may be closer to pre-exploitation levels than suggested by previous genetic assessments (*8, 9, 31*, Fig. 3, B), and the carrying capacity of the pre-exploitation oceans was possibly correspondingly lower. However, whale abundance estimated in this manner represents a temporal harmonic mean, and therefore necessarily reflects the actual abundance at a specific time period (*2*). Perhaps most importantly, our re-analysis demonstrated how sensitive this kind of inference is to changes in the underlying parameter values (*2*).

**Fig. 3.**
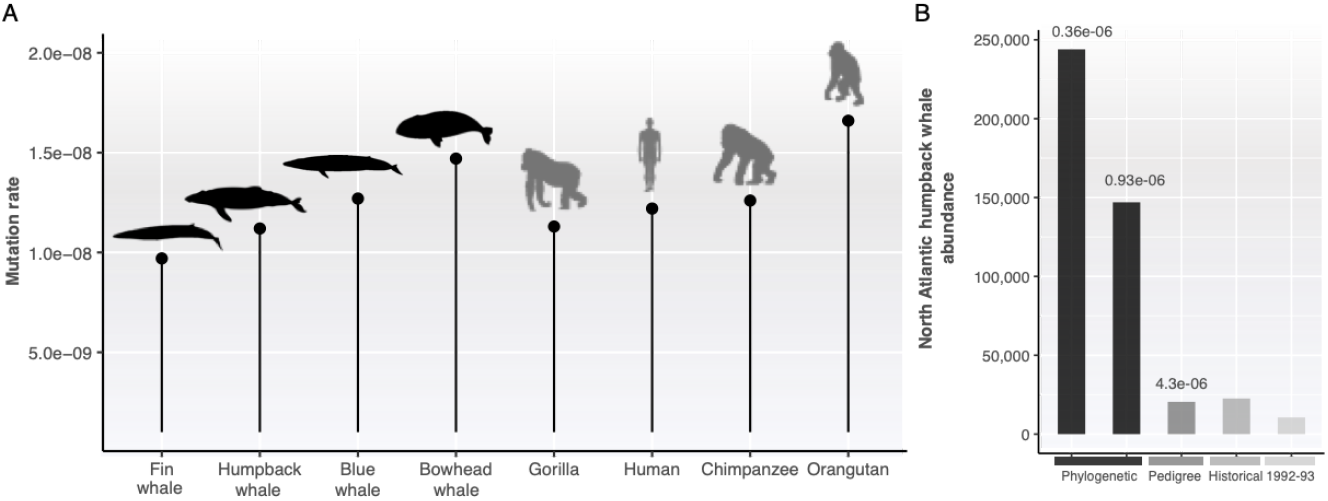
A) Pedigree-based estimates of genome mutation rates for baleen whales (this study) and large primates (table S1, *18, 32*). B) Estimates of contemporary and pre-exploitation abundance of North Atlantic humpback whales. Estimates based on *μ*_PHY_ were adjusted to match the generation time used in the original analysis and the annual rate of *μ*_PHY_ scaled to generations (18 years, *8, 9*). Historical abundances estimated through non-genetic means (*10*). Most recent published abundance estimate (*31*).

The rate and nature of *DNMs* is at the center of evolution itself and in many ecological and evolutionary inferences. Differences in *μ* among taxonomic groups have fostered a wide array of intriguing hypotheses aimed at understanding their underlying causes (*1*). However, there are technical caveats in the estimation of *μ*_PHY_ that render comparisons difficult because the observed variation might reflect biological or analytical differences, as well as incomplete records. This study shows that direct, pedigree-based estimation of *μ* in natural populations of non-model species is entirely feasible without any prior knowledge, even in comparatively inaccessible large mammals with exceptionally wide ranges and long generation times. The results of the pedigree-based approach to directly estimate *μ*_PED_ in baleen whales have notable implications for studies of human impacts on the ecosystem and evolutionary cancer biology. Given the relative ease of detecting parent-offspring trios in large samples and the low costs of whole genome sequencing, we foresee a bright future in obtaining pedigree-based estimates of *μ*_PED_ across a wide range of biodiversity. The pedigree-based approach generates directly comparable data, which will facilitate novel insights across a range of fundamental and applied aspects. Pedigree-based detection of hundreds to thousands of *DNMs* will also provide new and more detailed insights into the nature and distribution of *DNMs* thereby improving evolutionary and population genetic inference methods in general.

## Acknowledgements

We are indebted to the many staff who undertook the field and laboratory work over the years, and the long-term population research programs that made this work possible. We thank drs. Steve Beissinger and Oscar Gaggiotti for their valuable comments on this manuscript. The Center for Information Technology of the University of Groningen provided access and support to its Peregrine High Performance Computing cluster. We are grateful to Frédérique Lucas for providing whale drawings.

## Funding

M.SM was supported by a doctorate fellowship from the University of Groningen. P.J.P and M.B acknowledge the support over the years by the University of Copenhagen, University of California Berkeley and Stockholm University. Additional funding was provided by the Aage V. Jensen Foundation (P.J.P), US National Marine Fisheries Service (P.J.P, M.B), the International Whaling Commission Scientific Committee (M.B., P.J.P), WWF-DK (P.J.P.), the Commission for Scientific Research in Greenland (P.J.P), the Greenland Home Rule (P.J.P), the European Union’s Horizon 2020 Research and Innovation Programme under the Marie Skłodowska-Curie grant agreement no. 813383 (F.F., P.J.P.), and Consejo Nacional de Ciencia y Tecnología (V.RL.).

## Author contributions

M.SM., M.B., V.RL, J.R., and P.J.P. conceived and designed the study. J.R., R.S., C.R., and M.H-J. provided samples and field data. M.B. and M.SM. conducted the laboratory work. M.SM., V.RL. and F.F. performed the bioinformatic and data analyses supervised by P.J.P. M.SM., M.B., and P.J.P interpreted the results with contributions from all authors. M.SM. and P.J.P, wrote the manuscript with input from all authors. All authors approved the final version.

## Competing interests

The authors declare no competing interests.

## Supplementary Materials

### Materials and Methods

#### Samples

Skin biopsies were collected from free-ranging whales using a crossbow and modified arrows (*1*). The biopsies were stored in 6M NaCl with 25% dimethyl sulfoxide (*34*) at −80°C before DNA extraction. Total cell DNA was extracted by standard phenol/chloroform extractions as described by Russel and Sambrook (*35*) or using QIAGEN DNEasy^TM^ Blood and Tissue Kit (Qiagen Inc.) following the manufacturer’s instructions. Extracted DNA was stored in 1xTE buffer (10 mM Tris–HCl, 1 mM EDTA, pH 8.0) at −20°C.

### Nuclear mutation rate estimation

#### Paternity exclusion

Trios of whales were found by employing a paternity exclusion approach. Sex was determined by the differential amplification of the zinc finger coding regions, ZFX and ZFY(*36*) and by co-amplification of an SRY-specific region with one or more autosomal microsatellite loci (Bérubé, in prep). Samples were genotyped to up to 30-33 microsatellite markers (Bérubé, in prep). Every potential combination of a female with another whale was considered a putative mother-calf pair. These pairs were then compared with every male in the dataset. We sequenced the genome of all males that matched a mother-calf pair at every locus.

#### Genome sequencing

Whole-genome re-sequencing was carried out by the Beijing Genomic Institute. Pair-end (PE) sequencing, 100bp read-length, with free-PCR libraries, was performed on the BGISEQ500 platform for humpback, blue, and bowhead whale trios. Due to sample degradation, a modified protocol was employed for the bowhead whale samples in which DNA fragmentation was skipped. Fin whale samples were sequenced on a BGISEQ500 PE150 platform.

#### Genotype calling

The Bergeron et al. (*16*) pipeline was used to process the data and detect de novo mutations (*DNMs*) following the general guidelines described by Bergeron et al. (*17*). BWA-MEM (V0.7.17, *7*) was used to align the sequencing reads to the blue whale reference assembly (GCF_009873245.2, *8*) with the default options. PICARD (V2.25, *9*) was used to mark and remove duplicate reads. An initial set of variants was generated for recalibrating base quality scores. HaplotypeCaller and GenotypeGVCF were used to call variants. SNPs were filtered with GATK’s (v4.1.8, *10*) recommended hard filtering thresholds (QD > 2.0, FS < 60.0, MQ > 40.0, MQRankSum > −12.5, ReadPosRankSum > −8.0, SOR < 3.0). The remaining SNPs were used as “known” variants by BaseRecalibrator to recalibrate quality scores on the initial set of aligned reads. After recalibrating the quality scores, variants were called again with HaplotypeCaller in BP_RESOLUTION mode. CombineGVCFs was used to merge gVCF files per autosomal chromosome and GenomicsDBImport, samples per trio. Joint genotyping of the combined gVCF files was performed with GenotypeGVCF and the same hard site filters were applied to the resulting VCFs. Kinship coefficients were calculated employing VCFtools (V0.1.16, *11*) with the --relatedness2 parameter employing all biallelic sites along the autosomal genome. In all trios, the kinship coefficient between the parents and the calf was >0.2 confirming the relatedness relation inferred from the microsatellite markers.

#### Identification of de novo mutations

Using GATK’s VariantFiltration, mendelian violations were selected as potential *DNMs*. These are sites where the genotype of the offspring is not consistent with the genotypes of the parents based on Mendelian inheritance. Several filters were applied to the resulting mendelian violations to identify DNM candidates.

1. Both parents had to be homozygous for the reference allele with no reads assigned to the alternative allele (AD=0).
2. The number of reads for the alternative allele had to be between 30% and 70% of the total number of reads.
3. The read depth on each individual had to be between 0.5x-2x its average depth.
4. The genotype quality for a site had to be higher than 60 (GQ>60) on every individual of a trio.
5. At least one read of each strand had to support the alternative allele.

De novo mutation candidates that passed the previous filter were manually curated to detect false positives (fig. S4). Integrated Genomic Viewer was used to visualize realigned reads yielded by HaplotypeCaller. Candidate DNMs were considered false positives if the alternative allele on the offspring was supported just by reads on just one strand and there were multiple SNPs on the same reads not present in the parents. Although our pipeline resulted in a considerably higher number of putative *DNMs*, these were easy to identify and manually removed and it did not affect the performance of the pipeline (See Pipeline testing).

#### Phasing mutations

*DNMs* were phased by employing a read-based method described by Maretty et al. (*41*) This method employs reads that contain a *DNM* and at least another variable site that allows identifying the origin of the *DNM* if only one of the parents presents it.

#### Sanger sequencing DNMs

Primer-BLAST (*42*) was used to design primers to amplify and sequence a subset of *DNMs* detected in the blue whale trio and the humpback whale trio (table S4). The quality of the primers was further checked with AmplifX (v1.7, *14*). PCRs were carried out in 10 μl volumes; 1μl Taq Buffer (Tris-HCL 0.67M, MgCl_2_ 0.02M, (NH_4_)_2_SO_4_ 0.166M, β-mercaptoethanol 0.11M), 4μl dNTPs (2mM of each nucleotide), 1μl forward and reverse primers, 0.08μl Taq DNA polymerase, 1μl DNA template and 1.92μl of dubbed-distilled water. The PCR program was comprised of an initial denaturation at 92°C for two minutes, followed by 35 cycles of 15 seconds at 95°C, 15 seconds at 55°C and 30 seconds at 72°C. The program terminated with five minutes at 72°C. Amplification products were checked by electrophoresis through 2% Agarose gels in 1X TBE buffer (Tris-borate-EDTA) at 175V for 30 minutes, followed by staining with ethidium bromide and visualization under fluorescent light. PCR products were cleaned with Shrimp Alkaline Phosphatase and Exonuclease I, as described by Werle et al. (*44*). Cycle sequencing of cleaned PCR products was undertaken using the primers mentioned above and the BigDye ^®^ Terminator v3.1 Cycle Sequencing kit Applied Biosystems ™ Inc.) following the manufacturer’s protocol. The cycle sequencing products were purified by ethanol sodium precipitation. The order of sequencing fragments was resolved by electrophoresis on an ABI 3730 DNA Analyzer TM (Applied Biosystems Inc.).

#### Gene annotation and putative functional effect

*DNMs* were annotated and classified using SNPEFF (v4.3, *16*) to check the gene position and primary functional impact effect. This software uses a reference database to annotate mutations according to their respective genomic position and allocate an impact category based on functional impacts, such as amino acid changes. First, the SNPEFF –build function was employed to create a local database using the blue whale reference assembly gene transfer format (GTF) annotation file as a reference (NCBI accession number: GCF_009873245.2). VCFs containing *DNMs* were annotated and classified based on gene orthology using only canonical transcripts (-canon), and excluding downstream and upstream changes (-no-downstream-no-upstream). This filtering parameter avoids over-assignment of the putative impacts, leaving solely the most conservative putative impact effects. *DNMs* were then placed in one of each of the SNPEFF impact categories: *high* (e.g. loss of function mutations causing stop gain, stop loss, exon loss), *moderate* (e.g., missense with possible protein function disturbance, coding sequence mutations), *low* (non-detrimental for protein function, e.g, synonymous mutations), and *modifier* (non-impact variants, e.g, intergenic or intronic mutations). Any *DNM* presenting annotation warnings or errors were not considered and excluded from the analysis.

#### Estimating callable sites

To calculate the mutation rate per base pair, it was necessary to estimate the genome size in which *DNM* could be detected. For this step, the approach used by Bergeron et al was followed. Only sites where all individuals passed the following filters were considered; a) Hard site filters recommended by GATK (QD > 2.0, FS < 60.0, MQ > 40.0, MQRankSum > −12.5, ReadPosRankSum > −8.0, SOR < 3.0), b) both parents had to be homozygous to the reference allele with no reads assigned to the alternative alleles, c) read depth had to be 0.5x-2x the average read depth of the trio, d) the genotype quality scores had to be at least 60. A false-negative rate (FNR, α) due to the allelic balance filter was calculated; sites that were not called heterozygous on the calf despite both parents being homozygous for different alleles were counted and divided by the total amount of heterozygous sites. The mutation rate per generation was calculated employing the following equation (*16*); where *m* is the number of *DNM, β* the false positive rate (FPR), α the FNR and *C* the size of the callable genome.

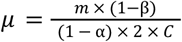

#### Pipeline testing

Bergeron and colleagues (*17*) analysed a trio of rhesus macaques employing five different bioinformatic pipelines. The same trio was analysed with our pipeline. Employing exactly the same parameters we used for the whales resulted in a mutation rate (0.38×10^-8^ substitutions per site per generation) below what other groups obtained in Bergeron et al. (2022). This consisted of 18*DNMs* detected with no false positives and 88% of the genome covered. We consider that the parameters of a pipeline should be adjusted based on the type of data employed. Bergeron et al. (2022) employed a trio sequenced at a considerable higher sequencing depth (~60X). Thus, we found that with our pipeline and high coverage data the AD=0 filter is too stringent. Allowing AD≤2 resulted in similar results as other groups (0.46×10^-8^-0.85×10^-8^) with a mutation rate of 0.62×10^-8^, 30 *DNMs* detected after manual curation (204 before), one false positive and 88% of the genome covered. As no false positives were detected by sequencing putative *DNMs*, the FPR obtained by analysing the macaque trio (*β =*0.033) was employed to estimate the mutation rates.

### Mitochondrial mutation rate estimation

#### Sequencing

Samples were genotyped at 19 microsatellite markers; AC087 (*46*) EV001, EV037, EV094, EV096 (*47*), GATA028, GATA053, GATA098, GATA417, TAA031 (*48*) GATA97408 (Bérubé, in prep) GT011 (*49*), GT015, GT023, GT101, GT195, GT211, GT271, GT575 (*50*) following the protocol described in (*51*). The sequence of the first 500 base pairs (bp) at the 5’ end of the mitochondrial control region was determined as described in Palsbøll et al. (*52*) using the primers BP16071R (*53*) and MT4F (*54*). Polymerase Chain Reaction (PCR, *25*) products were cleaned with Shrimp Alkaline Phosphatase and Exonuclease I, as described by Werle et al. (*44*). Cycle sequencing of cleaned PCR products was undertaken using the primers mentioned above and the BigDye ^®^ Terminator v3.1 Cycle Sequencing kit Applied Biosystems ™ Inc.) following the manufacturer’s protocol. The cycle sequencing products were purified by ethanol sodium precipitation. The order of sequencing fragments was resolved by electrophoresis on an ABI 3730 DNA Analyzer TM (Applied Biosystems Inc.).

#### Pedigree analysis

Through longitudinal studies, individual humpback whales were identified from their natural markings, including the pigmentation pattern of the ventral side of the fluke, the serrations along the trailing edge of the fluke and the shape and scarring of the dorsal fin (*56*). Mother-calf pairs were identified in the field based on their close, consistent association, stereotypical behaviors and size differential (*57*). Parentage analysis and pedigree estimation were carried out in the software FRANZ (*58*). The age at first possible reproduction was set at five years old for both sexes. The maximum number of candidates was set at 7,000 for fathers and 1,500 for mothers (Nmmax and Nfmax, respectively). Five chains were carried out for the simulated annealing with a maximum of 100,000,000 iterations. The Metropolis Hasting parameters consisted of ten chains with a burn-in of 2,000,000 followed by 40,000,000 and sampling every 200 iterations. FRANZ was run with different seeds ten times to estimate the consistency rate. Every maternal lineage present in the pedigree was extracted. A maternal lineage was defined as a founding mother -a mother without a known mother in the pedigree- and all her descendants through the maternal line.

#### Heteroplasmy discovery

PHFINDER (v1.0 *29*) was used to detect point heteroplasmy in the Sanger sequence electropherograms generated for samples in the maternal lineages. The first 500 bp of the mitochondrial control region of *M. novaeangliae*, started at position 15,490 and ended at 15,989 according to the mitogenome sequence NC_006927.1 (Genbank Reference Sequence, *30*). This sequence was used as reference for the alignment in PHFINDER. Heteroplasmies were sought in a 275bp segment that started at position 15,539 and ended at position 15,813 of the reference sequence. The heteroplasmy detection threshold was set at 15%, average base call quality 30, and secondary ratios 0.4. Lineages were removed when no electropherogram could be analyzed for the founding mother or in any individual in a first-generation. Whales for which no electropherogram could be analysed (low quality or incomplete sequence) were removed from the dataset. A detected heteroplasmy was considered to be present in the germ line and hence, transmittable to the offspring when the same heteroplasmy was detected in the founding mother and in at least one individual of the subsequent generations in the maternal line. For the purpose of this study, only maternal lineages in which a germ-line heteroplasmy was detected in the founding mother of a lineage were used for downstream analysis.

#### NGS sequencing

The mitochondrial genome was sequenced from skin samples of each individual in maternal lineages with a germ-line heteroplasmy to obtain a higher resolution of heteroplasmic frequencies. From several of these individuals, multiple skin samples were sequenced to estimate a measurement variance (see Bottleneck estimation). NGS library preparation was carried out following NEBNext Ultra II FS DNA Library Prep Kit for Illumina NEB #E7805) protocol, using NEBNext ^®^ Multiplex Oligos for Illumina ^®^ Dual Index Primers Set 1, NEB #7600). Default settings of the manufacturer’s protocol were followed during library preparation. The starting material was 390ng of purified genomic DNA in a final volume of 26μl. The selected size for the fragmentation stage was 150bp-350bp. During the size selection of adaptor-ligated DNA, an insert size distribution of 150bp-250bp was selected as the final library size distribution of 270bp-350bp. Hybridization of the prepared libraries was carried out using myBaits^®^ Mito kit Arbor Biosciences©) following the manufacturer’s protocol. This kit uses biotinylated RNA “baits” complementary to sequences of the humpback whale mitogenome (Genbank NC_006927.1), enriching the mitochondrial DNA in the library. The hybridization incubation was carried out for 43 hours. After the non-targeted DNA was washed away, the libraries were amplified through PCR for 18 cycles following the manufacturer’s protocol. The final dual-indexed library was diluted between 5-20 nM. Libraries sequencing was outsourced to The Hospital for Sick Children (Toronto, Canada) where paired-end Illumina next-generation sequencing (125bp) was carried out on the HiSeq2500 platform. Demultiplexing and adapter removal was performed at the sequencing site.

#### NGS data processing and heteroplasmy detection

The raw reads processing and calculation of minimum allele frequencies (MAF) for analysed heteroplasmies consisted of four stages.

##### Step 1 – Numt filtering

All raw reads were aligned with BOWTIE2 (v2.3.4.3, *31*) as “end to end” alignment employing the setting very_sensitive to the humpback whale nuclear assembly (Genbank GCA_004329385.1, *32*) supplemented with the mitochondrial control region (CR) with 250 extra base pairs at both ends to account for the mitogenome circularity (coordinates: 15234-16398, 1-250. Genbank NC_006927.1). SAMTOOLS (v1.9, *34*) was used to convert the resulting Sequence Alignment Map (SAM) file into a BAM file. After this, the BAM file was sorted and indexed (this process was repeated after each modification of a BAM file throughout the pipeline). From this BAM file, only paired reads that mapped in proper pair and primary alignment to the supplemented CR were kept for the downstream analysis. BEDTOOLS (v2.27.1, *35*) tool bamtofastq was used to convert the BAM file back into FASTQ format.

##### Step 2 – Consensus sequence

A consensus sequence was created for each sample. Reads obtained from Step 1 were realigned to the same CR (coordinates: 15234-16398, 1-250. Genbank NC_006927.1) with BOWTIE2 as “end to end” alignment employing the setting very_sensitive. SAMTOOLS was used to convert the resulting SAM file into a BAM file, removing some reads in the process (reads unmapped, mate unmapped, not primary alignments, reads failing platform, duplicates), retaining properly paired reads and filtering by a mapping quality of 20. Read groups were added to the BAM file using the PICARD (v2.18.5,*9*) tool AddOrReplaceReadGroups. The HaplotypeCaller tool from GATK (v4.1 *10*) was used (default settings) to generate a Variant Call Format (VCF) file (*40*). The VCF file was used by GATK’s tool FastaAlternateReferenceMaker to generate the consensus sequence of the sample. Finally, BLAST+ (v2.7.1) tool Dustmasker (*64*) was used to set in lowercase low complexity regions (i.e. regions containing simple sequence repeats) in the consensus FASTA sequence.

##### Step 3 – Alternative alleles read count

The second step generated a VCF file from which the MAF of potential heteroplasmies was obtained. For this, forward and reverse reads were aligned against the sample’s consensus reference with BOWTIE2 (very sensitive settings). SAMTOOLS was used to convert the resulting SAM file into a BAM file as in Step 1. The tool MarkDuplicates from PICARD was used to remove PCR/optical duplicates. A custom Python script was then used to filter from the SAM file reads that contained more than three mismatches; either INDELs or substitutions. After mismatch filtering, a second custom Python script was used to filter PCR/sequencing artefacts due to palindromic regions. For every read with mismatches, the script parsed the sequence of nucleotides from the mismatch position to the closest extreme of the read, either 5’ or 3’. Then, using Biopython v1.73 (*65*) the reverse complement of the selected sequence was generated and aligned to its immediate region in the reference sequence. If every position aligned perfectly, the read was discarded. After this custom filtering, the SAM file was checked for errors with CleanSam from PICARD. The SAM file was then converted again to a BAM file. FixMateInformation from PICARD was used to verify mate-pair information between read mates. BCFTOOLS (v1.9, *38*) commands mpileup (filtering by mapping and per base quality; settings -q 30 -Q 30 -C 50) and call (settings: --ploidy 1 -A -m) were used to obtain a VCF file from which the number of reads assigned to each alternative allele were parsed.

##### Step 4 – Heteroplasmy detection

Each position’s alleles were called based on the read counts extracted from the VCF files. The MAF was calculated as the percentage of reads that supported the second most common allele. For a position with multiple alleles to be considered heteroplasmic, we set a heteroplasmy detection limit similar to the one set by González et al. (*67*); MAF of 3% with read coverage depths of at least 500 and 1.5% MAF for depths of 3000x or more.

#### Bottleneck estimation

The effective size of the germ-line bottleneck that occurs during egg production was estimated following the model used by Millar et al. (*26*). This model is based on the assumption that when both variants of an heteroplasmy are neutral (i.e. under no selection), the inheritance will be based on a binomial distribution with parameters p, the mother’s MAF, and N, the germ-line bottleneck size in segregating units. Thus, the actual variance in MAF between mother and calf (from now on, genetic variance, σ^2^genetic) will be dependent on the aforementioned parameters as shown in equation (1).

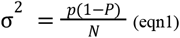

Millar and colleagues estimated the genetic variance as the raw variance (σ^2^raw) minus two times the heteroplasmy measurement variance (equation 3). In this equation, σ^2^raw is the squared difference between the maternal and the calf MAFs (equation 2) and σ^2^measure (measurement variance) is the uncertainty measuring the heteroplasmy proportions. The measurement variance is subtracted twice as two measurements are taken for each mother-calf pair.

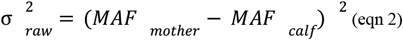

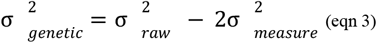

Transmission events where the MAF between mother and calf do not change would result in negative values of genetic variance when employing equation (3). To prevent this, we estimated the genetic variance by both adding and subtracting the measurement variance from every measurement (equation 4). This produced an upper and lower limit of the actual MAF value (Mu, Ml, maternal upper and lower MAF value respectively. Cu, Cl, calf upper and lower MAF value respectively). The measurement variance was calculated by obtaining the pooled variance from the MAFs from whales with a germ-line heteroplasmy in which more than one biopsy was sequenced (table S6). This accounted not only for technical uncertainties in the heteroplasmy measurement but also for other biological uncertainties, although with several limitations. A better approach to estimate the effects of non-germ-line bottlenecks in the measured heteroplasmy proportions would be to use samples from different tissues (i.e. skin and blood). This would not only take into account the effects of mitotic bottlenecks but also the developmental ones from embryonic tissue differentiation (*68, 69*). Obtaining different tissues from live whales was not feasible, thus assuming that the obtained measurement variation due to mitotic bottlenecks equals the overall variation (germ-lime, developmental and mitotic bottlenecks) was a necessary assumption. Several studies in other species (*69, 70*) have shown that the effect of the germ-line bottleneck is far more significant in determining heteroplasmy proportions. The raw variance between mother and calf was calculated for all four possible value combinations. The genetic variance for the mother-calf pair was set as the average of the raw variances (equation 4).

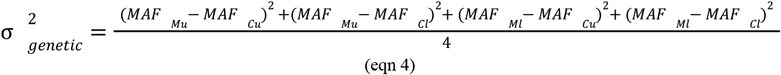

#### Mutation rate estimation

The mitochondrial control region germ-line mutation rate (μ), was estimated as described by Hendy et al. (*71*) and Millar et al. (*26*). They defined mutation rate as “the rate at which a base substitution is incorporated into all mitochondrial genomes of an individual”. Their model defines that new mutations, assuming neutrality and an equal probability to be transmitted, enter the germ line at rate α. Of these mutations, only 1/N are expected to go to fixation, meaning that μ=α/N. To be able to detect an heteroplasmy, the MAF has to be higher than a set threshold δ dependent on the detection method used. In this study, for Sanger sequencing chromatograms the detection limit was set to δ=0.15. The model assumes that given δ, most heteroplasmies are lost without being detected and most heteroplasmies that reach δ do not go to fixation. Thus, the proportion of sites with observed heteroplasmies (*β*) can be approximated as *β*= 2αln(1/δ −1). Being *β* the number of founding mothers with a detected germ-line heteroplasmy divided by the number of maternal lineages times the number of base pairs analysed. A simulation was used to correct the unequal number of whales and biopsies in heteroplasmic pedigrees. Only the first generation of each lineage was considered. For each simulation, one random sample from the founding mother and a sample from a random calf was chosen. Then it was checked whether using only those samples a germline heteroplasmy would have been detected. From 100 simulations, we obtained an average of 6 lineages in which germ line heteroplasmy would have been detected with equal sampling. Solving for α (μ=α/N) in *β*= 2αln(1/δ −1) one obtains equation (6) used to estimate the mutation rate per generation.

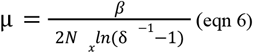

#### Population size estimation

The resulting mutation rate was used to estimate the long-term effective population size of North Atlantic humpback whales as done by Roman and Palumbi (*72*). With the exception of the mutation rate, all parameters were kept the same for the purpose of comparison. We employed equation *θ*=*Ne_f_μ*, being *θ* the genetic diversity of the population, *Ne_f_* the effective population size of females and μ, our estimate of the mutation rate per generation. We used the estimates of *θ*=0.022 calculated in their study as the section of the control region used in both studies largely overlaps (210bp). Total census size was calculated employing a conversion factor of eight; *Ne_f_* was converted to total Ne by multiplying by two, considering a 1:1 sex ratio (*73*). Ne was then converted to the total number of breeding adults employing a ratio of two (*74*). Finally, to account for juveniles we multiplied again by two (*72, 75*).

**Fig. S1.**
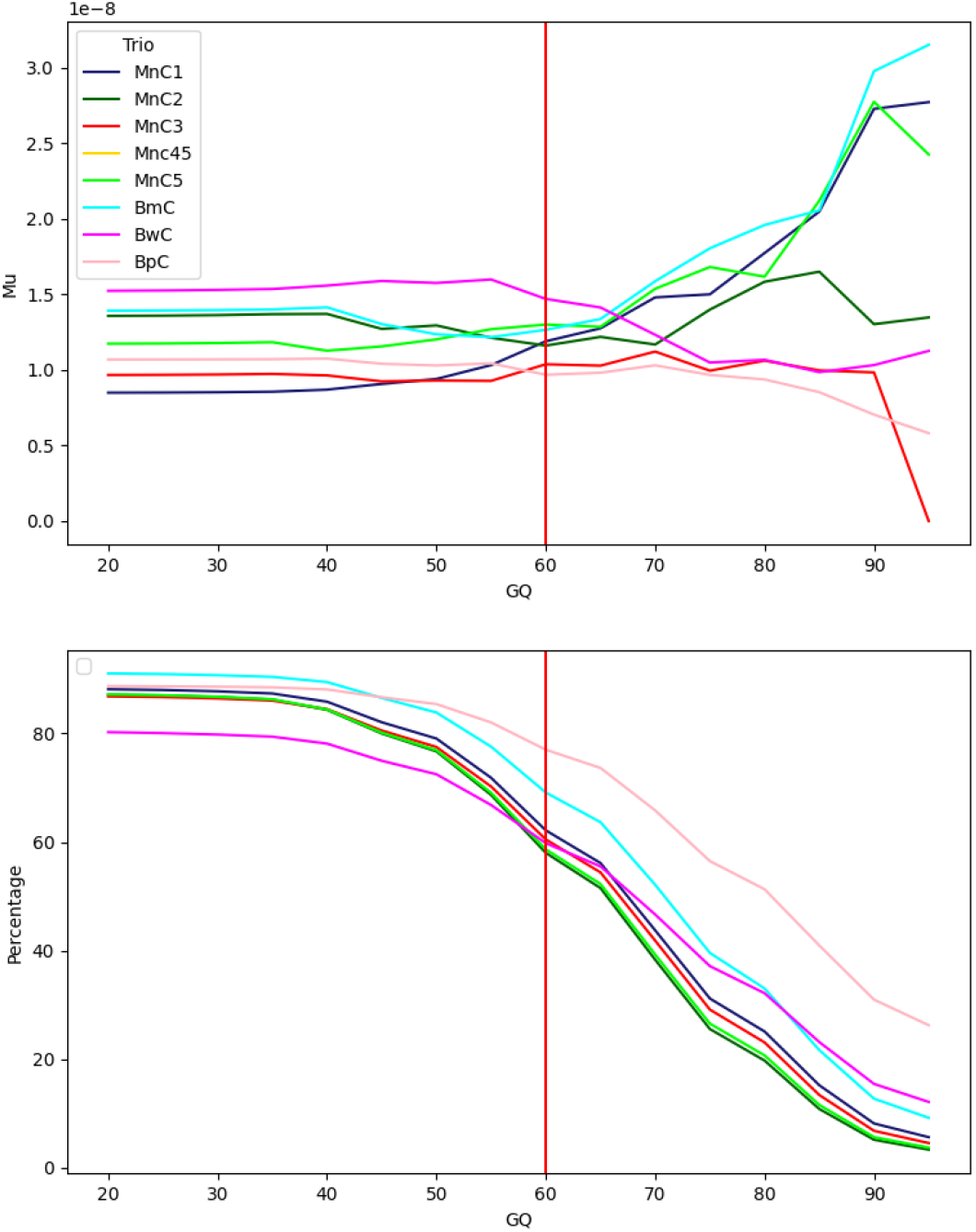
Changes in the mutation rate (top) and percentage of the callable genome (bottom) with different genotype quality (GQ) thresholds.

**Fig. S2.**
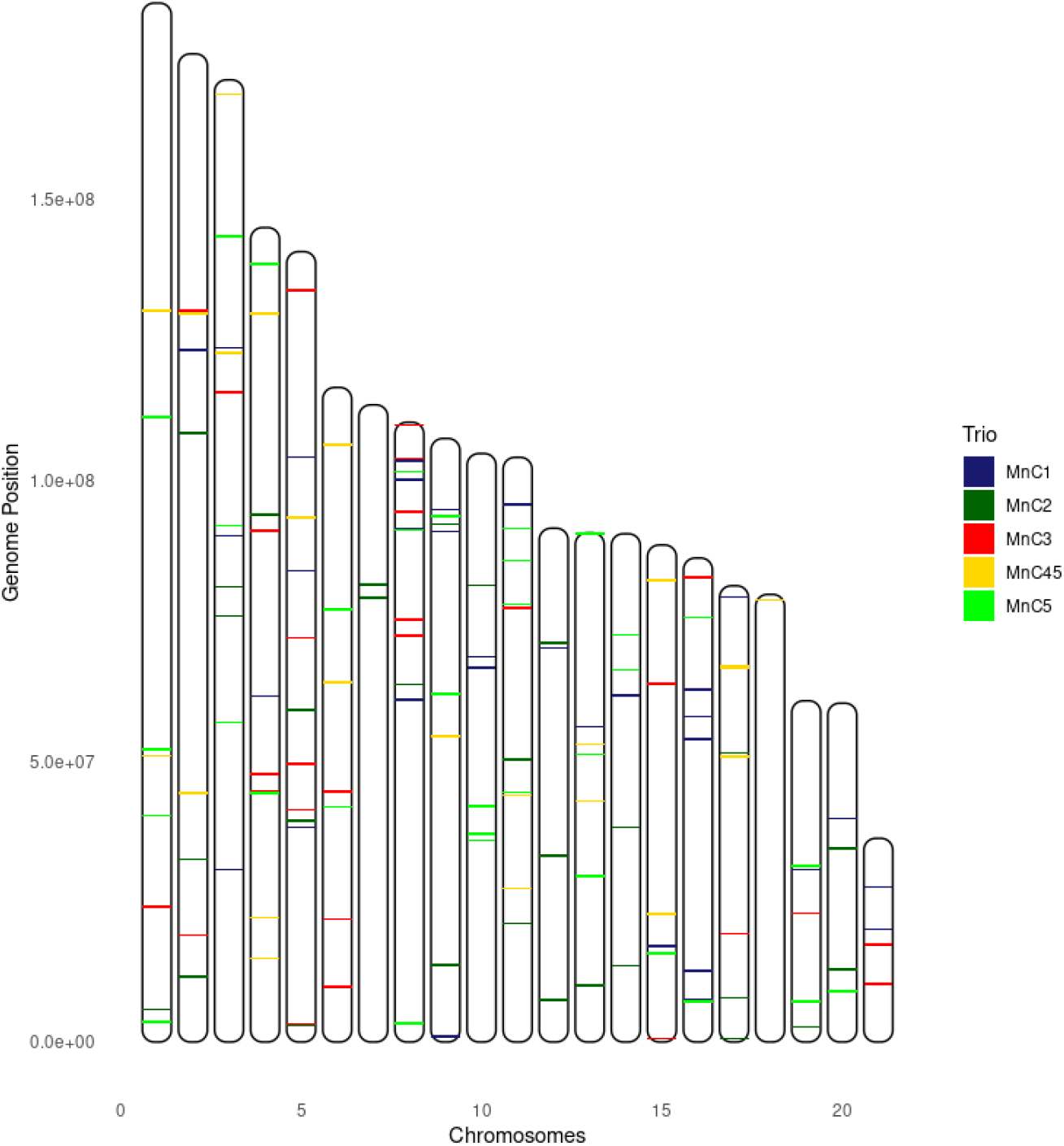
Location of *DNMs* on all humpback whale trios along the chromosomes.

**Fig. S3.**
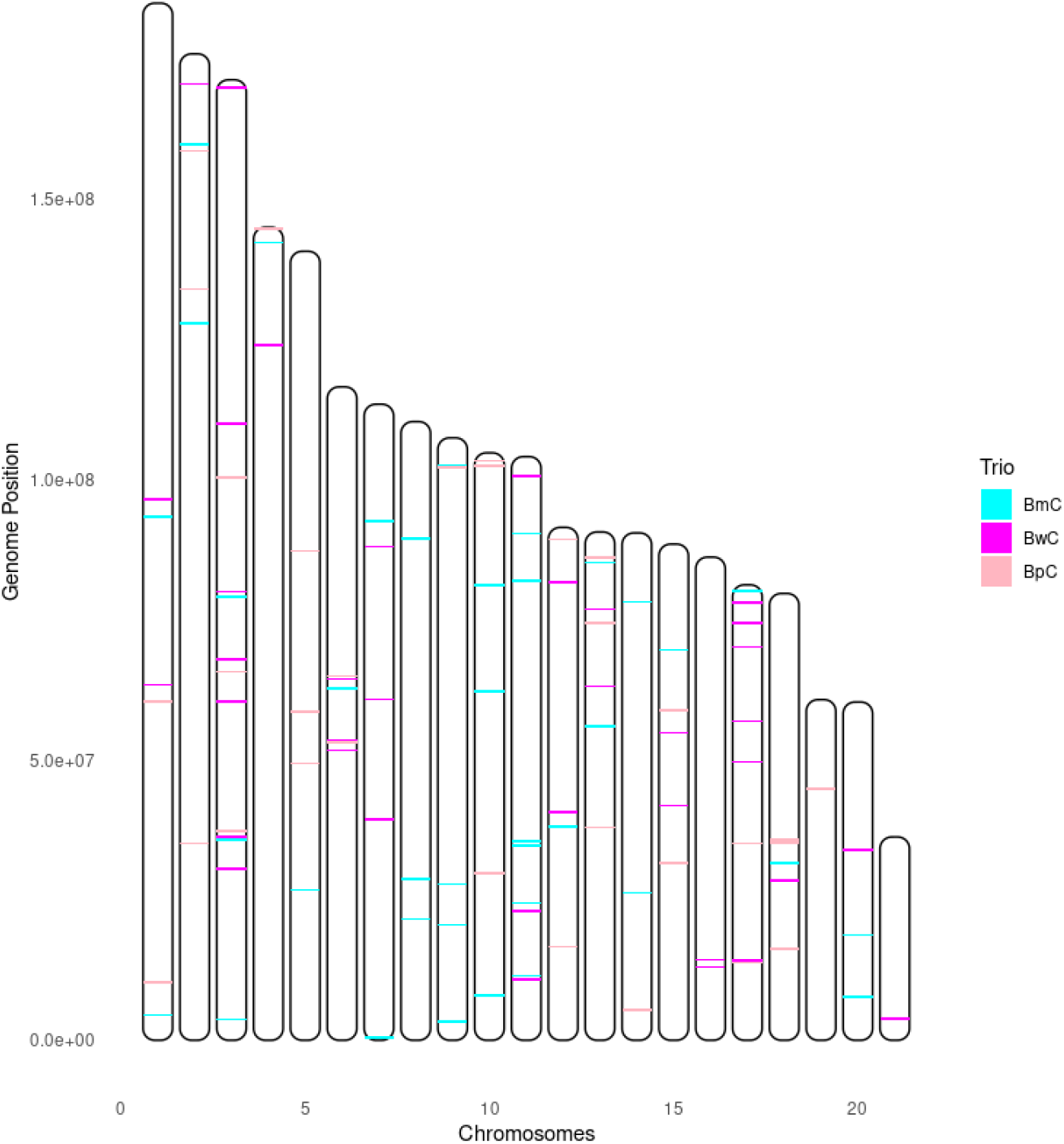
Location of *DNMs* on the blue, fin and bowhead whale trios along the chromosomes.

**Fig. S4.**
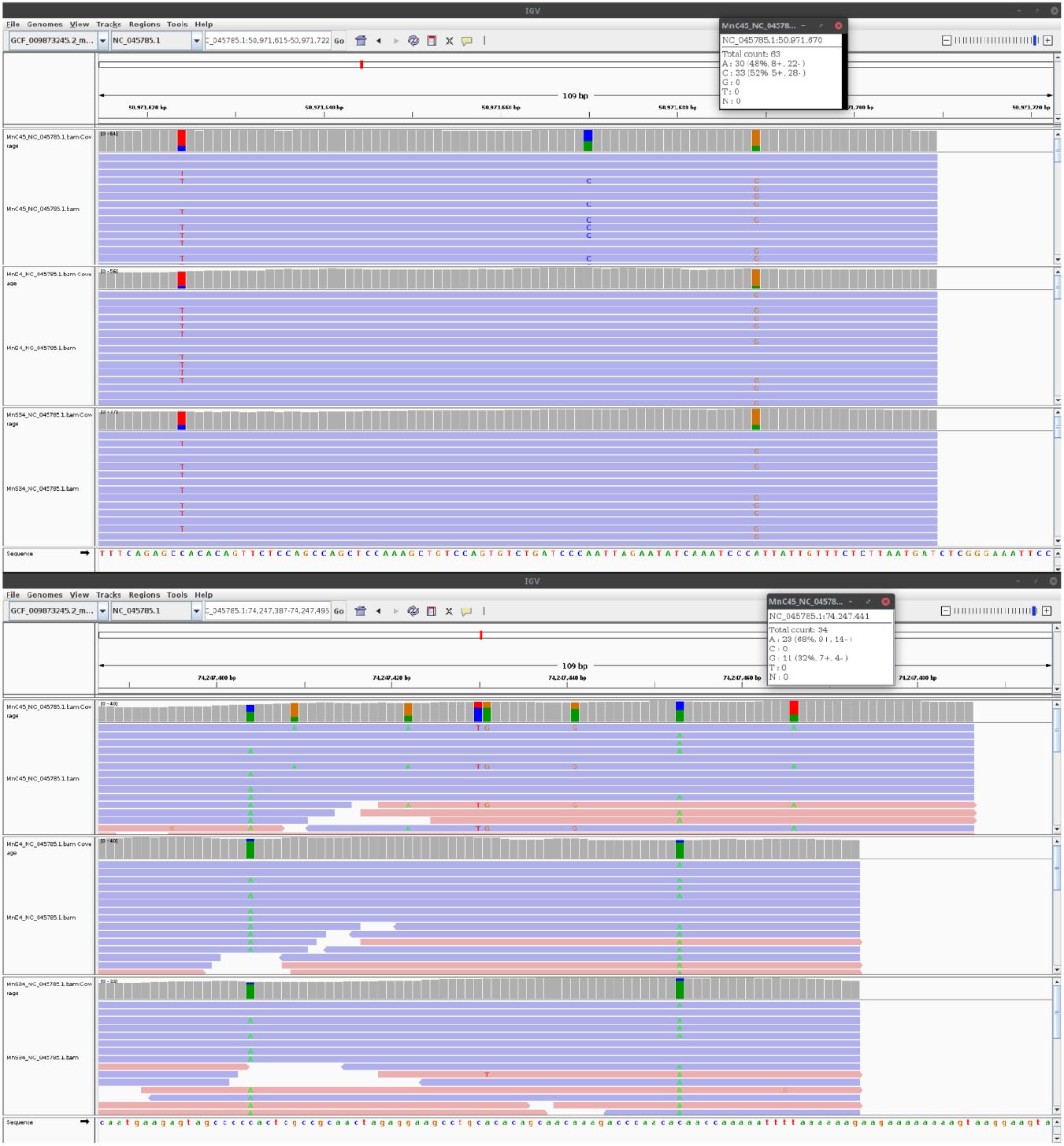
Example of manual curation of *DNMs* on IGV viewer. Top image shows a putative *DNM* that passed curation. From top to bottom, calf, mother and father. Bottom image shows a putative *DNM* that did not pass manual curation.

**Fig. S5.**
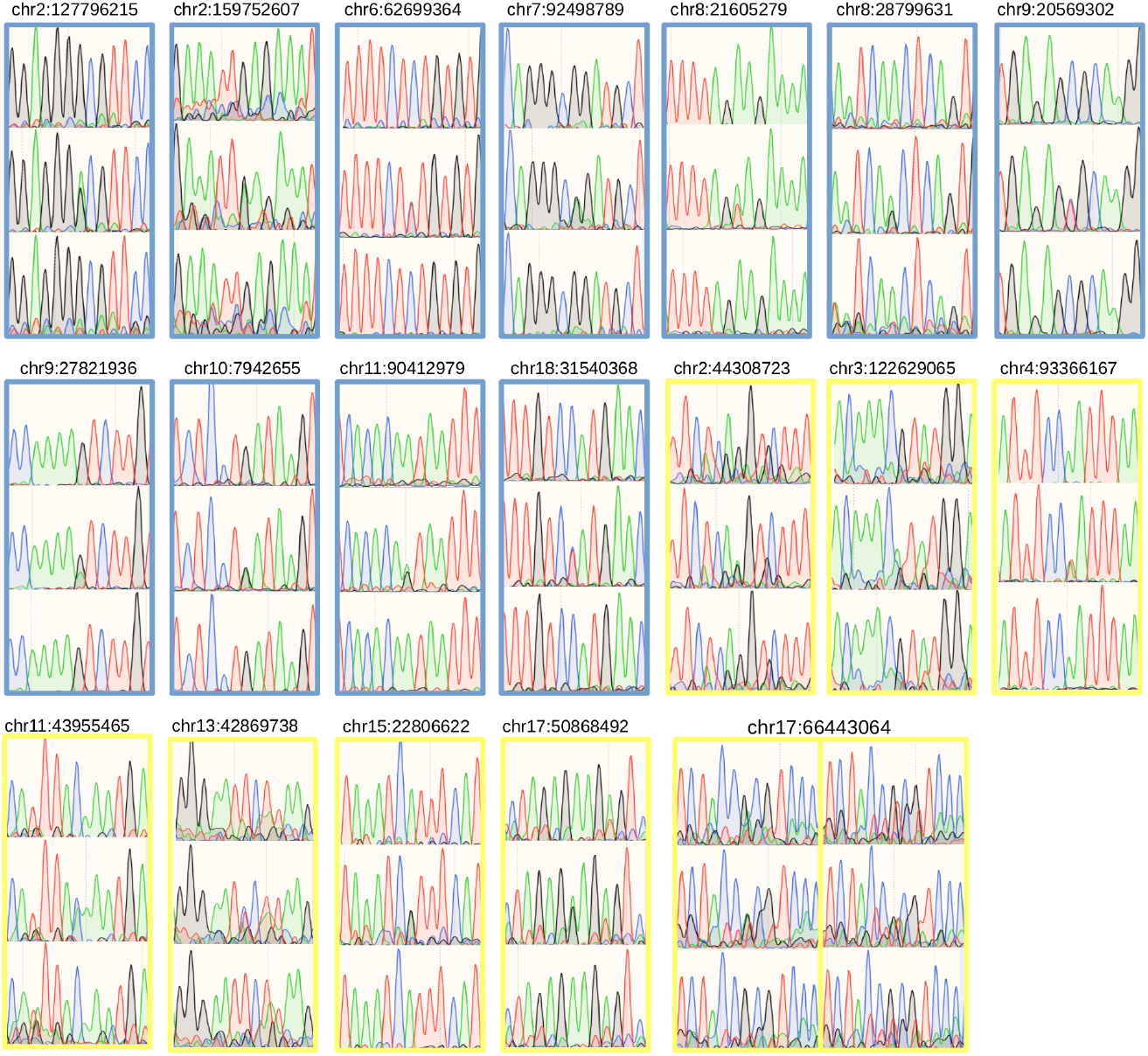
Sanger sequencing chromatograms of 19 *DNM* that successfully amplified and sequenced. Blue rectangles indicate a *DNM* from the blue whale trio (BmC), in yellow humpback whale trio (MnC45). Top chromatogram, mother, middle calf, bottom father. *DNM* mutation in the middle position of each chromatogram.

**Fig. S6.**
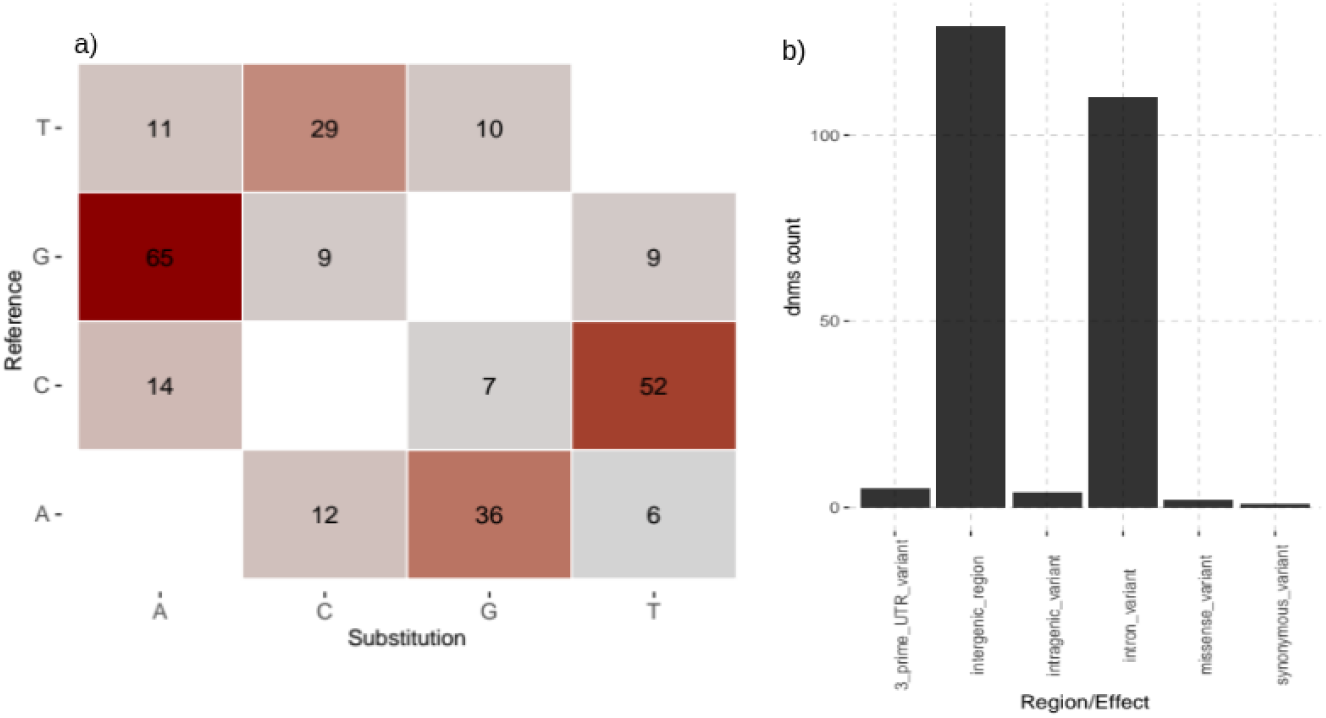
Characterization of *DNM*. a) Type of *DNM*. b) Region and effect of *DNMs*.

**Fig. S7.**
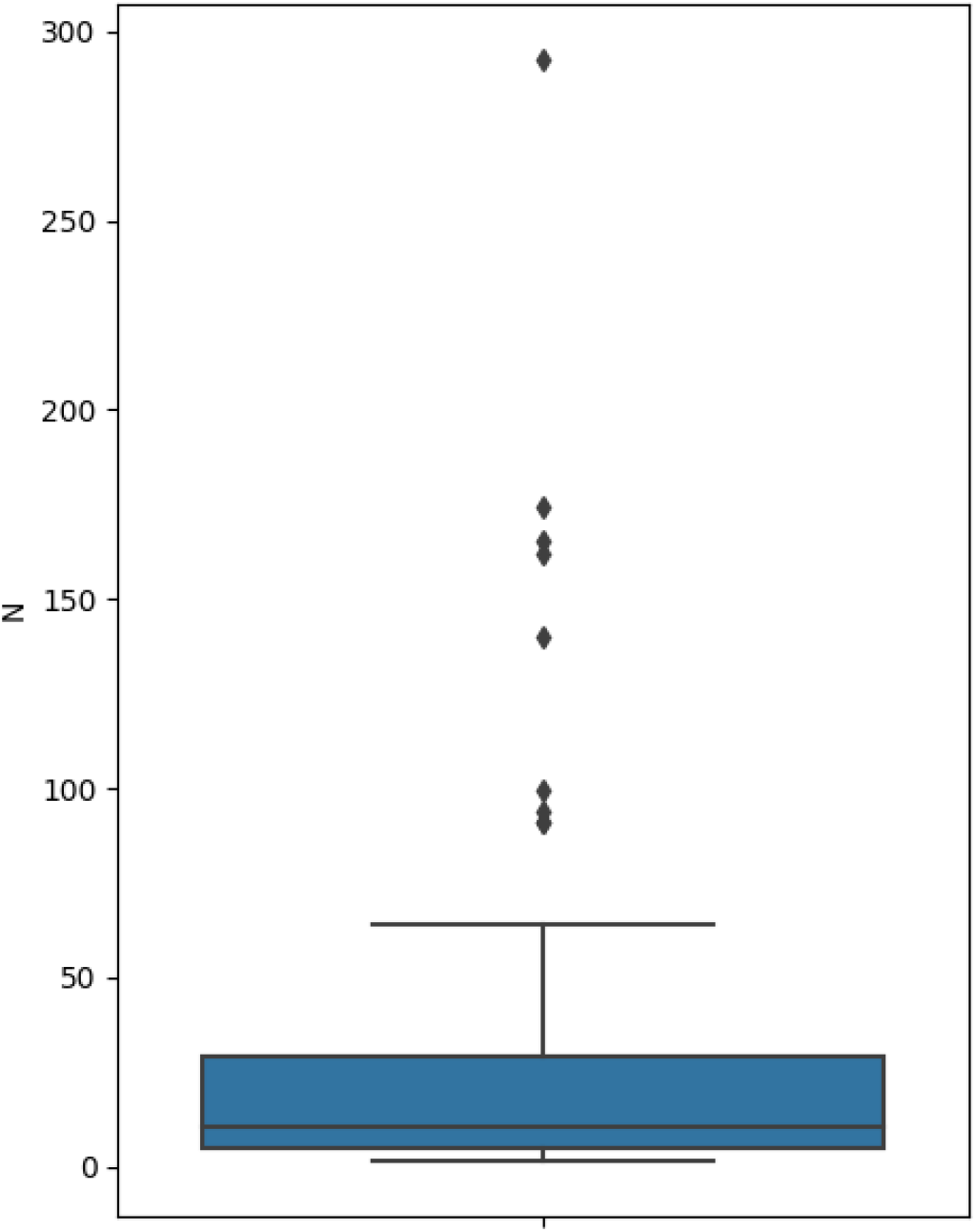
Estimated number of segregating units (N) estimated from the change of heteroplasmic frequencies.

**Table S1.** Direct estimates of nuclear mutation rates on vertebrate species.

| Species | Mu per generation | Reference |
| --- | --- | --- |
| Herring (*Clupea harengus*) | 0.2E-08 | (76) |
| Cichlids (*Astatotilapia calliptera*, *Aulonocara stuartgranti*, and *Lethrinops lethrinus*) | 0.35E-08 | (77) |
| Pig (*Sus scrofa*) | 0.36E-08 | (78) |
| Mouse (*Mus musculus*) | 0.39E-08 | (79) |
| Marmoset (*Callithrix jacchus*) | 0.43E-08 | (80) |
| Gray wolf (*Canis lupus*) | 0.45E-08 | (81) |
| Collared flycatcher (*Ficedula albicollis*) | 0.46E-08 | (13) |
| Baboon (*Papio anubis*) | 0.57E-08 | (82) |
| Platypus (*Ornithorhynchus anatinus*) | 0.7E-08 | (11) |
| Rhesus macaque (*Macaca mulatta*) | 0.77E-08 | (16) |
| Owl monkey (*Aotus nancymae*) | 0.81E-08 | (83) |
| Brown bear (*Ursus arctos horribilis*) | 0.84E-08 | (84) |
| Cat (*Felis catus*) | 0.86E-08 | (85) |
| Green monkey (*Chlorocebus sabaeus*) | 0.94E-08 | (86) |
| Gorilla (*Gorilla gorilla*) | 1.13E-08 | (87) |
| Cattle (*Bos Taurus*) | 1.17E-08 | (88) |
| Human (*Homo sapiens*) | 1.22E-08 | (32) |
| Chimpanzee (*Pan troglodytes*) | 1.26E-08 | (87) |
| Gray mouse lemur (*Microcebus murinus*) | 1.52E-08 | (89) |
| Orangutan (*Pongo abelii*) | 1.66E-08 | (87) |

**Table S2.**
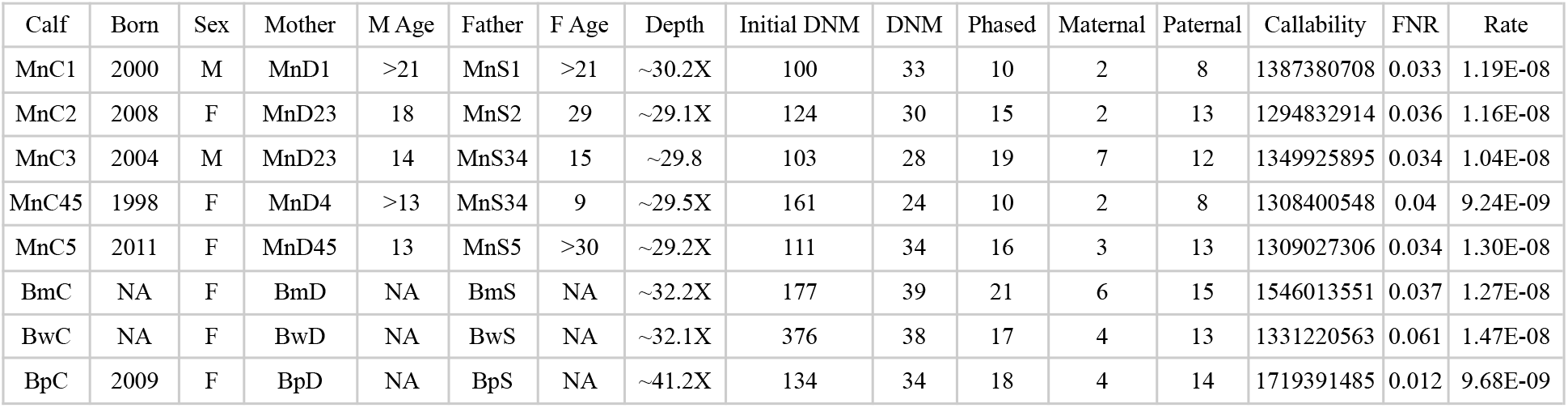
Information about each trio: year calf was born, calf sex, age or minimum age of parents when calf was born, average depth of each trio after filters, number of *DNM* obtained with the pipeline and number of *DNM* that passed manual curation, phased mutation to each parent, number of bases of the autosomal genome that passed all filters, false negative rate and mutation rate.

| Calf | Born | Sex | Mother | M Age | Father | F Age | Depth | Initial DNM | DNM | Phased | Maternal | Paternal | Callability | FNR | Rate |
| --- | --- | --- | --- | --- | --- | --- | --- | --- | --- | --- | --- | --- | --- | --- | --- |
| MnC1 | 2000 | M | MnD1 | >21 | MnS1 | >21 | ~30.2X | 100 | 33 | 10 | 2 | 8 | 1387380708 | 0.033 | 1.19E-08 |
| MnC2 | 2008 | F | MnD23 | 18 | MnS2 | 29 | ~29.1X | 124 | 30 | 15 | 2 | 13 | 1294832914 | 0.036 | 1.16E-08 |
| MnC3 | 2004 | M | MnD23 | 14 | MnS34 | 15 | ~29.8 | 103 | 28 | 19 | 7 | 12 | 1349925895 | 0.034 | 1.04E-08 |
| MnC45 | 1998 | F | MnD4 | >13 | MnS34 | 9 | ~29.5X | 161 | 24 | 10 | 2 | 8 | 1308400548 | 0.04 | 9.24E-09 |
| MnC5 | 2011 | F | MnD45 | 13 | MnS5 | >30 | ~29.2X | 111 | 34 | 16 | 3 | 13 | 1309027306 | 0.034 | 1.30E-08 |
| BmC | NA | F | BmD | NA | BmS | NA | ~32.2X | 177 | 39 | 21 | 6 | 15 | 1546013551 | 0.037 | 1.27E-08 |
| BwC | NA | F | BwD | NA | BwS | NA | ~32.1X | 376 | 38 | 17 | 4 | 13 | 1331220563 | 0.061 | 1.47E-08 |
| BpC | 2009 | F | BpD | NA | BpS | NA | ~41.2X | 134 | 34 | 18 | 4 | 14 | 1719391485 | 0.012 | 9.68E-09 |

**Table S3.** List of positions of all *DNMs* and whether they were unphased (U), phased to the father (P) or to the mother (M). In bold *DNM* found in MnC45 that were transmitted to MnC5.

| Calf | Chromosome | *DNMs* | Phase |
| --- | --- | --- | --- |
| MnC1 | Chr2 | 123250494 | U |
| MnC1 | Chr3 | 30712312 | P |
| MnC1 | Chr3 | 90142215 | U |
| MnC1 | Chr3 | 123579014 | U |
| MnC1 | Chr4 | 61539962 | U |
| MnC1 | Chr5 | 38273433 | P |
| MnC1 | Chr5 | 83913722 | U |
| MnC1 | Chr5 | 104123224 | P |
| MnC1 | Chr8 | 60934463 | U |
| MnC1 | Chr8 | 91400989 | U |
| MnC1 | Chr8 | 100087168 | P |
| MnC1 | Chr8 | 103424558 | U |
| MnC1 | Chr9 | 955366 | U |
| MnC1 | Chr9 | 90922533 | U |
| MnC1 | Chr9 | 94805965 | U |
| MnC1 | Chr10 | 66614021 | P |
| MnC1 | Chr10 | 68624075 | U |
| MnC1 | Chr11 | 95663131 | U |
| MnC1 | Chr12 | 70155619 | U |
| MnC1 | Chr13 | 56170965 | M |
| MnC1 | Chr14 | 61639631 | U |
| MnC1 | Chr15 | 17035534 | P |
| MnC1 | Chr15 | 17071708 | P |
| MnC1 | Chr16 | 7560394 | U |
| MnC1 | Chr16 | 12720908 | U |
| MnC1 | Chr16 | 53960431 | U |
| MnC1 | Chr16 | 57914348 | M |
| MnC1 | Chr16 | 62677306 | U |
| MnC1 | Chr17 | 79175402 | U |
| MnC1 | Chr19 | 30665231 | P |
| MnC1 | Chr20 | 39797956 | U |
| MnC1 | Chr21 | 20104100 | U |
| MnC1 | Chr21 | 27560283 | U |
| MnC2 | Chr1 | 5749459 | U |
| MnC2 | Chr2 | 11560971 | U |
| MnC2 | Chr2 | 32464180 | M |
| MnC2 | Chr2 | 108458519 | U |
| MnC2 | Chr3 | 75839400 | P |
| MnC2 | Chr3 | 80997127 | P |
| MnC2 | Chr4 | 93900090 | U |
| MnC2 | Chr5 | 3032521 | P |
| MnC2 | Chr5 | 39415348 | P |
| MnC2 | Chr5 | 59095879 | P |
| MnC2 | Chr7 | 79068637 | P |
| MnC2 | Chr7 | 81447750 | U |
| MnC2 | Chr8 | 63626960 | P |
| MnC2 | Chr9 | 13673951 | P |
| MnC2 | Chr9 | 92153049 | U |
| MnC2 | Chr10 | 81317192 | P |
| MnC2 | Chr11 | 21100371 | U |
| MnC2 | Chr11 | 50308875 | U |
| MnC2 | Chr12 | 7522864 | P |
| MnC2 | Chr12 | 33131010 | U |
| MnC2 | Chr12 | 71075631 | U |
| MnC2 | Chr13 | 10116027 | M |
| MnC2 | Chr14 | 13556870 | U |
| MnC2 | Chr14 | 38202285 | P |
| MnC2 | Chr17 | 576379 | U |
| MnC2 | Chr17 | 7872693 | U |
| MnC2 | Chr17 | 51447384 | P |
| MnC2 | Chr19 | 2645963 | U |
| MnC2 | Chr20 | 12895998 | P |
| MnC2 | Chr20 | 34394901 | U |
| MnC3 | Chr1 | 24060116 | P |
| MnC3 | Chr2 | 19076584 | U |
| MnC3 | Chr2 | 130210842 | U |
| MnC3 | Chr3 | 115614823 | U |
| MnC3 | Chr4 | 44718329 | P |
| MnC3 | Chr4 | 47756745 | P |
| MnC3 | Chr4 | 91063666 | P |
| MnC3 | Chr5 | 3200933 | P |
| MnC3 | Chr5 | 41325633 | M |
| MnC3 | Chr5 | 49443178 | M |
| MnC3 | Chr5 | 71904681 | M |
| MnC3 | Chr5 | 133763983 | P |
| MnC3 | Chr6 | 9753815 | U |
| MnC3 | Chr6 | 21902221 | P |
| MnC3 | Chr6 | 44613027 | P |
| MnC3 | Chr8 | 72313906 | U |
| MnC3 | Chr8 | 75254610 | P |
| MnC3 | Chr8 | 94380212 | M |
| MnC3 | Chr8 | 103890829 | M |
| MnC3 | Chr8 | 109814839 | U |
| MnC3 | Chr11 | 77329600 | P |
| MnC3 | Chr15 | 661033 | P |
| MnC3 | Chr15 | 63816311 | U |
| MnC3 | Chr16 | 82780118 | U |
| MnC3 | Chr17 | 19228213 | M |
| MnC3 | Chr19 | 22937366 | M |
| MnC3 | Chr21 | 10326195 | P |
| MnC3 | Chr21 | 17327609 | U |
| **MnC45** | **Chr1** | **50971670** | **P** |
| MnC45 | Chr1 | 130183287 | U |
| MnC45 | Chr2 | 44308723 | P |
| **MnC45** | **Chr2** | **129665965** | **U** |
| MnC45 | Chr3 | 122629065 | M |
| MnC45 | Chr3 | 168759839 | P |
| MnC45 | Chr4 | 14824695 | U |
| MnC45 | Chr4 | 22179901 | P |
| MnC45 | Chr4 | 129644441 | U |
| MnC45 | Chr5 | 93366167 | U |
| **MnC45** | **Chr6** | **64101747** | **P** |
| **MnC45** | **Chr6** | **106275881** | **U** |
| **MnC45** | **Chr9** | **54369065** | **U** |
| **MnC45** | **Chr9** | **93605498** | **U** |
| MnC45 | Chr11 | 27329095 | U |
| MnC45 | Chr11 | 43955465 | U |
| MnC45 | Chr13 | 42869738 | U |
| MnC45 | Chr13 | 52987561 | P |
| MnC45 | Chr15 | 22806622 | U |
| **MnC45** | **Chr15** | **82155882** | **P** |
| MnC45 | Chr17 | 50868492 | M |
| MnC45 | Chr17 | 66443064 | P |
| MnC45 | Chr17 | 66889389 | U |
| **MnC45** | **Chr18** | **78679788** | **U** |
| MnC5 | Chr1 | 3552682 | U |
| MnC5 | Chr1 | 40316187 | P |
| MnC5 | Chr1 | 52132963 | M |
| MnC5 | Chr1 | 111233370 | U |
| MnC5 | Chr3 | 56853017 | U |
| MnC5 | Chr3 | 91961399 | P |
| MnC5 | Chr3 | 143473699 | U |
| MnC5 | Chr4 | 44253217 | U |
| MnC5 | Chr4 | 138520467 | P |
| MnC5 | Chr6 | 41870066 | P |
| MnC5 | Chr6 | 77006757 | U |
| MnC5 | Chr8 | 3332313 | P |
| MnC5 | Chr8 | 91119236 | U |
| MnC5 | Chr8 | 101568353 | U |
| MnC5 | Chr9 | 61887573 | U |
| MnC5 | Chr9 | 93567799 | M |
| MnC5 | Chr10 | 35869682 | P |
| MnC5 | Chr10 | 37099324 | P |
| MnC5 | Chr10 | 41933963 | P |
| MnC5 | Chr11 | 44491167 | P |
| MnC5 | Chr11 | 77856270 | P |
| MnC5 | Chr11 | 85717474 | U |
| MnC5 | Chr11 | 91355529 | U |
| MnC5 | Chr13 | 29542476 | U |
| MnC5 | Chr13 | 51216018 | U |
| MnC5 | Chr13 | 90469329 | U |
| MnC5 | Chr14 | 66225375 | P |
| MnC5 | Chr14 | 72490633 | U |
| MnC5 | Chr15 | 15711117 | P |
| MnC5 | Chr16 | 7183804 | U |
| MnC5 | Chr16 | 75635651 | M |
| MnC5 | Chr19 | 7274992 | U |
| MnC5 | Chr19 | 31372998 | U |
| MnC5 | Chr20 | 9062327 | P |
| BmC | Chr1 | 4483519 | P |
| BmC | Chr1 | 93286434 | U |
| BmC | Chr2 | 127796215 | U |
| BmC | Chr2 | 159752607 | U |
| BmC | Chr3 | 3745675 | P |
| BmC | Chr3 | per785862 | P |
| BmC | Chr3 | 79074147 | P |
| BmC | Chr4 | 142304870 | M |
| BmC | Chr5 | 26858356 | P |
| BmC | Chr6 | 62699364 | P |
| BmC | Chr7 | 416662 | M |
| BmC | Chr7 | 92498789 | U |
| BmC | Chr8 | 21605279 | P |
| BmC | Chr8 | 28799631 | U |
| BmC | Chr8 | 89517243 | P |
| BmC | Chr9 | 3276241 | U |
| BmC | Chr9 | 20569302 | U |
| BmC | Chr9 | 27821936 | M |
| BmC | Chr9 | 102425941 | U |
| BmC | Chr10 | 7942655 | U |
| BmC | Chr10 | 62157582 | M |
| BmC | Chr10 | 81124648 | U |
| BmC | Chr11 | 10958728 | U |
| BmC | Chr11 | 11541947 | U |
| BmC | Chr11 | 24513042 | P |
| BmC | Chr11 | 34690236 | M |
| BmC | Chr11 | 35486234 | U |
| BmC | Chr11 | 81977232 | U |
| BmC | Chr11 | 90412979 | P |
| BmC | Chr12 | 38099499 | M |
| BmC | Chr13 | 56038527 | U |
| BmC | Chr13 | 85137147 | P |
| BmC | Chr14 | 26290112 | U |
| BmC | Chr14 | 78201420 | P |
| BmC | Chr15 | 69637287 | P |
| BmC | Chr17 | 80049486 | P |
| BmC | Chr18 | 31540368 | U |
| BmC | Chr20 | 7772222 | U |
| BmC | Chr20 | 18796296 | P |
| BpC | Chr1 | 10376411 | P |
| BpC | Chr1 | 60377696 | U |
| BpC | Chr2 | 35090542 | U |
| BpC | Chr2 | 133891768 | P |
| BpC | Chr2 | 158549327 | M |
| BpC | Chr3 | 37295543 | P |
| BpC | Chr3 | 65755303 | P |
| BpC | Chr3 | 100324656 | P |
| BpC | Chr4 | 144717008 | P |
| BpC | Chr5 | 49324046 | U |
| BpC | Chr5 | 58594696 | U |
| BpC | Chr5 | 87237324 | P |
| BpC | Chr6 | 53122206 | P |
| BpC | Chr6 | 64951470 | M |
| BpC | Chr6 | 64967041 | M |
| BpC | Chr9 | 102170648 | U |
| BpC | Chr10 | 29736182 | U |
| BpC | Chr10 | 102497190 | U |
| BpC | Chr10 | 103384078 | M |
| BpC | Chr12 | 16694364 | U |
| BpC | Chr12 | 89280044 | U |
| BpC | Chr13 | 37929915 | U |
| BpC | Chr13 | 74347019 | P |
| BpC | Chr13 | 86102353 | P |
| BpC | Chr14 | 5381311 | U |
| BpC | Chr15 | 31624787 | U |
| BpC | Chr15 | 58852704 | U |
| BpC | Chr17 | 14032573 | P |
| BpC | Chr17 | 35150861 | P |
| BpC | Chr18 | 16273753 | P |
| BpC | Chr18 | 35214863 | U |
| BpC | Chr18 | 35720207 | U |
| BpC | Chr18 | 35876726 | U |
| BpC | Chr19 | 44874727 | P |
| BwC | Chr1 | 63375732 | U |
| BwC | Chr1 | 96507115 | U |
| BwC | Chr2 | 170542042 | P |
| BwC | Chr3 | 30506100 | M |
| BwC | Chr3 | 36265009 | P |
| BwC | Chr3 | 60419107 | P |
| BwC | Chr3 | 67948875 | U |
| BwC | Chr3 | 79975192 | P |
| BwC | Chr3 | 109932469 | U |
| BwC | Chr3 | 169811934 | U |
| BwC | Chr4 | 124026351 | P |
| BwC | Chr6 | 51708370 | U |
| BwC | Chr6 | 53555190 | U |
| BwC | Chr6 | 64458460 | U |
| BwC | Chr7 | 39418501 | U |
| BwC | Chr7 | 60745229 | U |
| BwC | Chr7 | 88046196 | M |
| BwC | Chr11 | 10763065 | P |
| BwC | Chr11 | 10985568 | P |
| BwC | Chr11 | 23000153 | P |
| BwC | Chr11 | 100612370 | P |
| BwC | Chr12 | 40697849 | U |
| BwC | Chr12 | 81720861 | P |
| BwC | Chr13 | 63132061 | U |
| BwC | Chr13 | 76875822 | U |
| BwC | Chr15 | 41791794 | U |
| BwC | Chr15 | 54862159 | M |
| BwC | Chr16 | 13039351 | P |
| BwC | Chr16 | 14298429 | U |
| BwC | Chr17 | 14264147 | U |
| BwC | Chr17 | 49666977 | M |
| BwC | Chr17 | 56858682 | P |
| BwC | Chr17 | 70169596 | U |
| BwC | Chr17 | 74382084 | U |
| BwC | Chr17 | 78102952 | U |
| BwC | Chr18 | 28474071 | P |
| BwC | Chr20 | 33963838 | U |
| BwC | Chr21 | 3833003 | U |

**Table S4.**
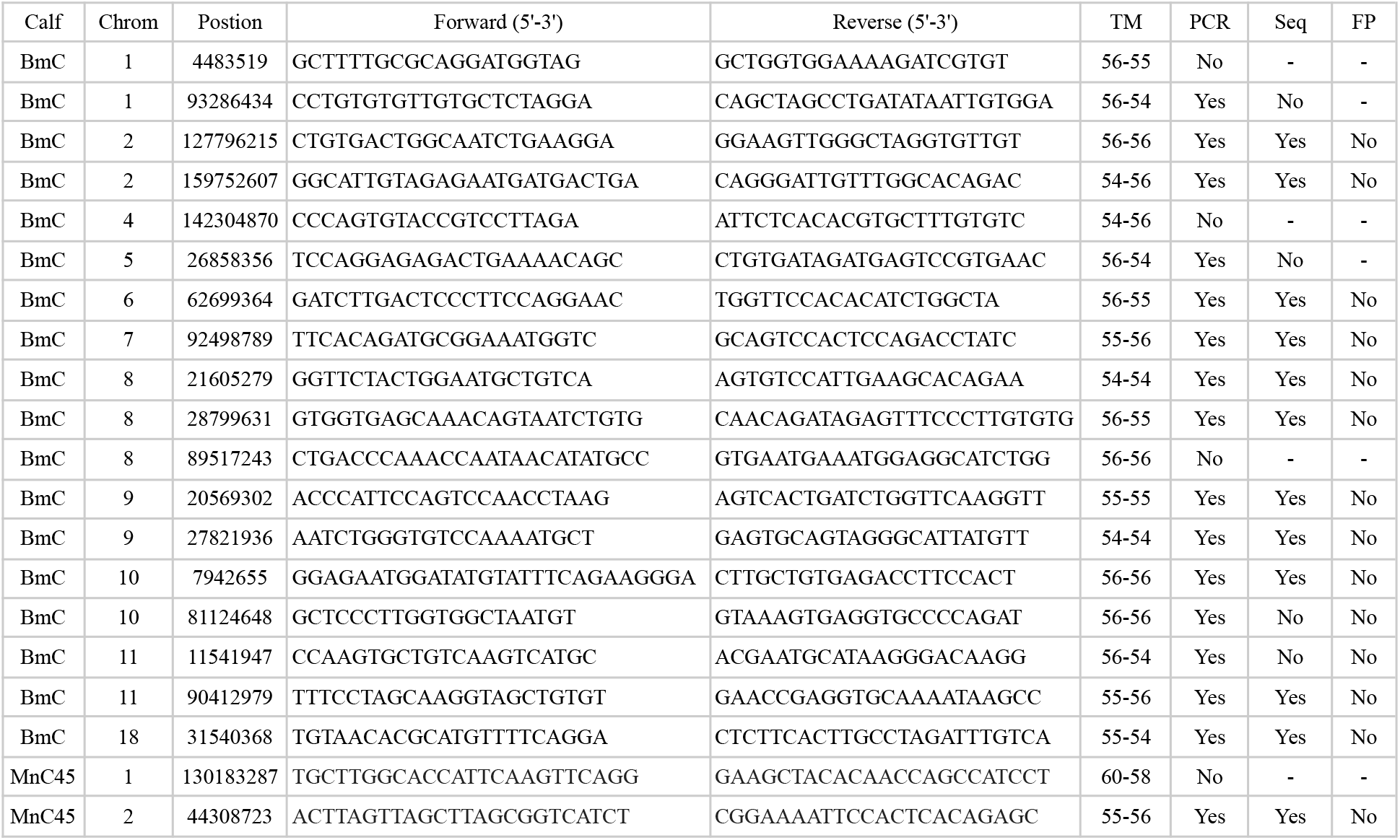
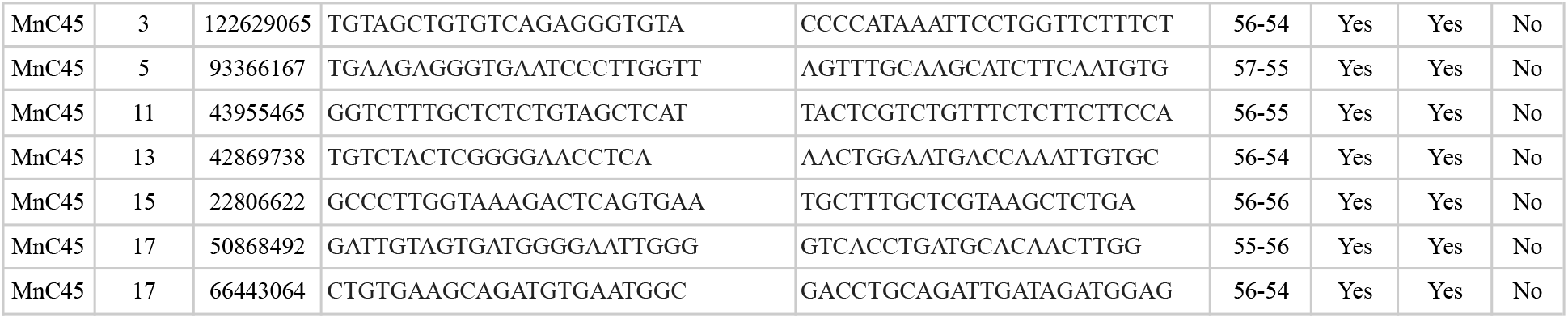
Primers designed for PCR validation of *DNMs*. TM, annealing temperature. PCR, whether all samples werere amplified correctly and specifically. Seq, whether Sanger sequencing produced clean chromatograms. FP, false positive.

**Table S5.** Heteroplasmic lineages detected, number of individuals, generations and main and secondary haplotypes (based on the most supported haplotype present the founding mother).

| Lineage | Position HP | Individuals | Generations | Main | Secondary |
| --- | --- | --- | --- | --- | --- |
| L11 | 258 | 6 | 2 | HapH | HapC |
| L14 | 82 | 16 | 3 | HapC | Novel |
| L47 | 258 | 3 | 2 | HapC | HapH |
| L67 | 235 | 4 | 2 | HapB | Novel |
| L76 | 82 | 7 | 3 | HapN | HapG |
| L97 | 82 | 3 | 2 | HapB | HapT |
| L105 | 258 | 9 | 3 | HapH | HapC |
| L115 | 82 | 4 | 2 | HapT | HapB |
| L127 | 82 | 5 | 2 | HapN | HapG |

**Table S6.** Heteroplasmy frequencies: position, mother, bases, number of reads assigned to each allele, heteroplasmic ratio, number of segregation units estimated (N) and estimated mutation rate.

| Position | Mother | HP | Highest | Lowest | Ratio | Calf | HP | Highest | Lowest | Ratio | N | Mu |
| --- | --- | --- | --- | --- | --- | --- | --- | --- | --- | --- | --- | --- |
| 82 | L127 | T/C | 5214 | 1926 | 27 | L127_1 | T/C | 2913 | 625 | 17.7 | 21 | 2.13E-06 |
| 82 | L127 | T/C | 5214 | 1926 | 27 | L127_2 | T/C | 3099 | 483 | 13.5 | 10 | 4.29E-06 |
| 82 | L127 | T/C | 5214 | 1926 | 27 | L127_4 | C/T | 624 | 523 | 45.6* | 3 | 1.72E-05 |
| 258 | L47 | G/A | 2850 | 1611 | 36.1 | L47_1 | G/A | 841 | 530 | 38.7 | 162 | 2.76E-07 |
| 258 | L47 | G/A | 2850 | 1611 | 36.1 | L47_2 | G/A | 4925 | 896 | 15.4 | 5 | 8.43E-06 |
| 235 | L67 | A/G | 8657 | 3119 | 26.5 | L67_1 | A/G | 1774 | 263 | 12.9 | 10 | 4.41E-06 |
| 235 | L67 | A/G | 8657 | 3119 | 26.5 | L67_2 | A/G | 5719 | 504 | 8.1 | 6 | 7.92E-06 |
| 235 | L67 | A/G | 8657 | 3119 | 26.5 | L67_3 | A/G | 6667 | 750 | 10.1 | 7 | 6.33E-06 |
| 82 | L76 | T/C | 8686 | 2404 | 21.7 | L76_5 | T/C | 6145 | 8 | 0 | 4 | 1.14E-05 |
| 82 | L76 | T/C | 8686 | 2404 | 21.7 | L76_1 | T/C | 1264 | 721 | 36.3 | 8 | 5.79E-06 |
| 82 | L76 | T/C | 8686 | 2404 | 21.7 | L76_2 | T/C | 8119 | 526 | 6.1 | 7 | 6.59E-06 |
| 82 | L76 | T/C | 8686 | 2404 | 21.7 | L76_4 | T/C | 4933 | 328 | 6.2 | 7 | 6.50E-06 |
| 82 | L76 | T/C | 8686 | 2404 | 21.7 | L76_3 | T/C | 1949 | 644 | 24.8 | 99 | 4.49E-07 |
| 82 | L76_2 | T/C | 8119 | 526 | 6.1 | L76_2.1 | T/C | 2609 | 204 | 7.3 | 64 | 6.96E-07 |
| 337 | L97 | C/T | 3426 | 1713 | 33.3 | L97_1 | T/C | 3616 | 2439 | 40.3* | 3 | 1.41E-05 |
| 337 | L97 | C/T | 3426 | 1713 | 33.3 | L97_2 | C/T | 6654 | 3278 | 33 | 293 | 1.52E-07 |
| 258 | L11 | A/G | 2175 | 1431 | 39.7 | L11_1 | G/A | 5839 | 2173 | 27.1* | 2 | 2.07E-05 |
| 258 | L11 | A/G | 2175 | 1431 | 39.7 | L11_2 | A/G | 1935 | 1413 | 42.2 | 174 | 2.56E-07 |
| 258 | L11 | A/G | 2175 | 1431 | 39.7 | L11_3 | G/A | 4900 | 2898 | 37.1* | 4 | 1.02E-05 |
| 258 | L11 | A/G | 2175 | 1431 | 39.7 | L11_4 | G/A | 4804 | 2610 | 35.2* | 4 | 1.19E-05 |
| 258 | L11 | A/G | 2175 | 1431 | 39.7 | L11_5 | A/G | 1996 | 495 | 19.8 | 6 | 7.52E-06 |
| 82 | L14 | C/T | 466 | 52 | 10 | L14_6 | C/T | 2275 | 577 | 20.2 | 8 | 5.53E-06 |
| 82 | L14 | C/T | 466 | 52 | 10 | L14_7 | C/T | 4032 | 174 | 4.1 | 21 | 2.10E-06 |
| 82 | L14 | C/T | 466 | 52 | 10 | L14_2 | C/T | 4656 | 769 | 14.2 | 36 | 1.25E-06 |
| 82 | L14 | C/T | 466 | 52 | 10 | L14_3 | C/T | 3027 | 141 | 4.5 | 24 | 1.87E-06 |
| 82 | L14 | C/T | 466 | 52 | 10 | L14_4 | C/T | 5720 | 2724 | 32.3 | 2 | 2.50E-05 |
| 82 | L14 | C/T | 466 | 52 | 10 | L14_5 | C/T | 703 | 3 | 0 | 10 | 4.28E-06 |
| 82 | L14 | C/T | 466 | 52 | 10 | L14_1 | C/T | 11774 | 16 | 0 | 10 | 4.28E-06 |
| 82 | L14_6 | C/T | 2275 | 577 | 20.2 | L14_6.1 | C/T | 5276 | 19 | 0 | 4 | 1.04E-05 |
| 82 | L14_6 | C/T | 2275 | 577 | 20.2 | L14_6.7 | C/T | 2468 | 12 | 0 | 4 | 1.04E-05 |
| 82 | L14_6 | C/T | 2275 | 577 | 20.2 | L14_6.3 | C/T | 5025 | 115 | 18.2 | 140 | 3.18E-07 |
| 82 | L14_6 | C/T | 2275 | 577 | 20.2 | L14_6.4 | C/T | 6041 | 1239 | 17 | 91 | 4.91E-07 |
| 82 | L14_6 | C/T | 2275 | 577 | 20.2 | L14_6.8 | C/T | 1824 | 66 | 3.5 | 6 | 7.92E-06 |
| 82 | L14_6 | C/T | 2275 | 577 | 20.2 | L14_6.6 | C/T | 3215 | 1999 | 38.3 | 5 | 9.27E-06 |
| 82 | L14_6 | C/T | 2275 | 577 | 20.2 | L14_6.2 | C/T | 3755 | 449 | 10.7 | 16 | 2.70E-06 |
| 82 | L14_6 | C/T | 2275 | 577 | 20.2 | L14_6.5 | C/T | 5173 | 1435 | 21.7 | 165 | 2.70E-07 |
| 258 | L105 | A/G | 6007 | 3261 | 35.2 | L105_3 | A/G | 2354 | 1057 | 31 | 91 | 4.91E-07 |
| 258 | L105 | A/G | 6007 | 3261 | 35.2 | L105_1 | A/G | 5856 | 2118 | 26.6 | 28 | 1.59E-06 |
| 258 | L105 | A/G | 6007 | 3261 | 35.2 | L105_2 | A/G | 9755 | 4413 | 31.1 | 94 | 4.75E-07 |
| 258 | L105 | A/G | 6007 | 3261 | 35.2 | L105_4 | A/G | 3319 | 2670 | 44.6 | 24 | 1.87E-06 |
| 258 | L105_1 | A/G | 5856 | 2118 | 26.6 | L105_1.1 | A/G | 7338 | 3996 | 35.2 | 24 | 1.86E-06 |
| 258 | L105_1 | A/G | 5856 | 2118 | 26.6 | L105_1.2 | A/G | 3877 | 2354 | 37.8 | 15 | 3.04E-06 |
| 258 | L105_1 | A/G | 5856 | 2118 | 26.6 | L105_1.3 | A/G | 5125 | 463 | 8.3 | 6 | 7.82E-06 |
| 258 | L105_1 | A/G | 5856 | 2118 | 26.6 | L105_1.4 | A/G | 2742 | 1465 | 34.8 | 26 | 1.71E-06 |
| 82 | L115 | T/C | 5240 | 2643 | 33.5 | L115_2 | C/T | 3728 | 1472 | 28.3 | 2 | 2.94E-05 |
| 82 | L115 | T/C | 5240 | 2643 | 33.5 | L115_3 | T/C | 12615 | 463 | 3.5 | 2 | 1.82E-05 |
| 82 | L115 | T/C | 5240 | 2643 | 33.5 | L115_1 | T/C | 1006 | 340 | 25.3 | 30 | 1.50E-06 |

\*Transmission in which the allele supported by the highest number of reads flipped to the alternate allele in the calf.

**Table S7.** Heteroplasmy frequencies estimated for multiple biopsies of three individuals and the inferred pooled measurement error of the estimations.

| Individual | Sample | Ratio |
| --- | --- | --- |
| L105_1 | MN0976 | 28.3 |
| L105_1 | MN0977 | 22.7 |
| L105_1 | MN0978 | 26.6 |
| L105 | MN0974 | 36.7 |
| L105 | MN0975 | 34.7 |
| L105 | MN1126 | 35.2 |
| L76 | MN0802 | 21.7 |
| L76 | MN0803 | 22.1 |
| Pooled measurement error |  | 0.019 |

## Notes

### Competing Interest Statement

The authors have declared no competing interest.

